# Asrij/OCIAD1 depletion reduces inflammatory microglial activation and ameliorates Aβ pathology in an Alzheimer’s disease mouse model

**DOI:** 10.1101/2024.08.15.608066

**Authors:** Prathamesh Dongre, Madhu Ramesh, Thimmaiah Govindaraju, Maneesha S. Inamdar

## Abstract

**Background:** Alzheimer’s disease (AD) is a neurodegenerative disorder characterized by the accumulation of amyloid-beta (Aβ) plaques and neurofibrillary tangles, neuroinflammation, and glial activation. Asrij/OCIAD1 (Ovarian Carcinoma Immunoreactive Antigen Domain containing protein 1) is an AD-associated factor. Increased Asrij levels in the brains of AD patients and mouse models are linked to the severity of neurodegeneration. However, the contribution of Asrij to AD progression and whether reducing Asrij levels is sufficient to mitigate Aβ pathology *in vivo* is unclear.

**Methods:** To explore the impact of Asrij on AD pathology, we deleted *asrij* in the *APP/PS1* mouse model of AD and analyzed the effects on AD hallmarks. We used the Morris water maze and open field test to assess behavioral performance. Using immunohistochemistry and biochemical analyses, we evaluated Aβ plaque load, neuronal and synaptic damage, and gliosis. Further, we utilized confocal microscopy imaging, flow cytometry and RNA sequencing analysis to comprehensively investigate changes in microglial responses to Aβ pathology upon Asrij depletion.

**Results:** Asrij depletion ameliorates cognitive impairments, Aβ deposition, neuronal and synaptic damage, and reactive astrogliosis in the AD mouse. Notably, Asrij-deficient microglia exhibit reduced plaque-associated proliferation and decreased phagocytic activation. Transcriptomic analyses of AD microglia reveal upregulation of energy metabolism pathways and downregulation of innate immunity and inflammatory pathways upon Asrij depletion. Mechanistically, loss of Asrij increases mitochondrial activity and impedes the acquisition of a pro-inflammatory disease-associated microglia (DAM) state. Reduced levels of pro-inflammatory cytokines and decreased STAT3 and NF-κB activation indicate protective changes in AD microglia. Taken together, our results suggest that increased Asrij levels reported in AD, may suppress microglial metabolic activity and promote inflammatory microglial activation, thereby exacerbating AD pathology.

**Conclusions:** In summary, we show that Asrij depletion ameliorates Aβ pathology, neuronal and synaptic damage, gliosis, and improves behavioral performance in *APP/PS1* mice. This supports that Asrij exacerbates the AD pathology. Mechanistically, Asrij is critical for the development of DAM and promotes neuroinflammatory signaling activation in microglia, thus restricting neuroprotective microglial responses. Hence, reducing Asrij in this context may help retard AD. Our work positions Asrij as a critical molecular regulator that links microglial dysfunction to AD pathogenesis.

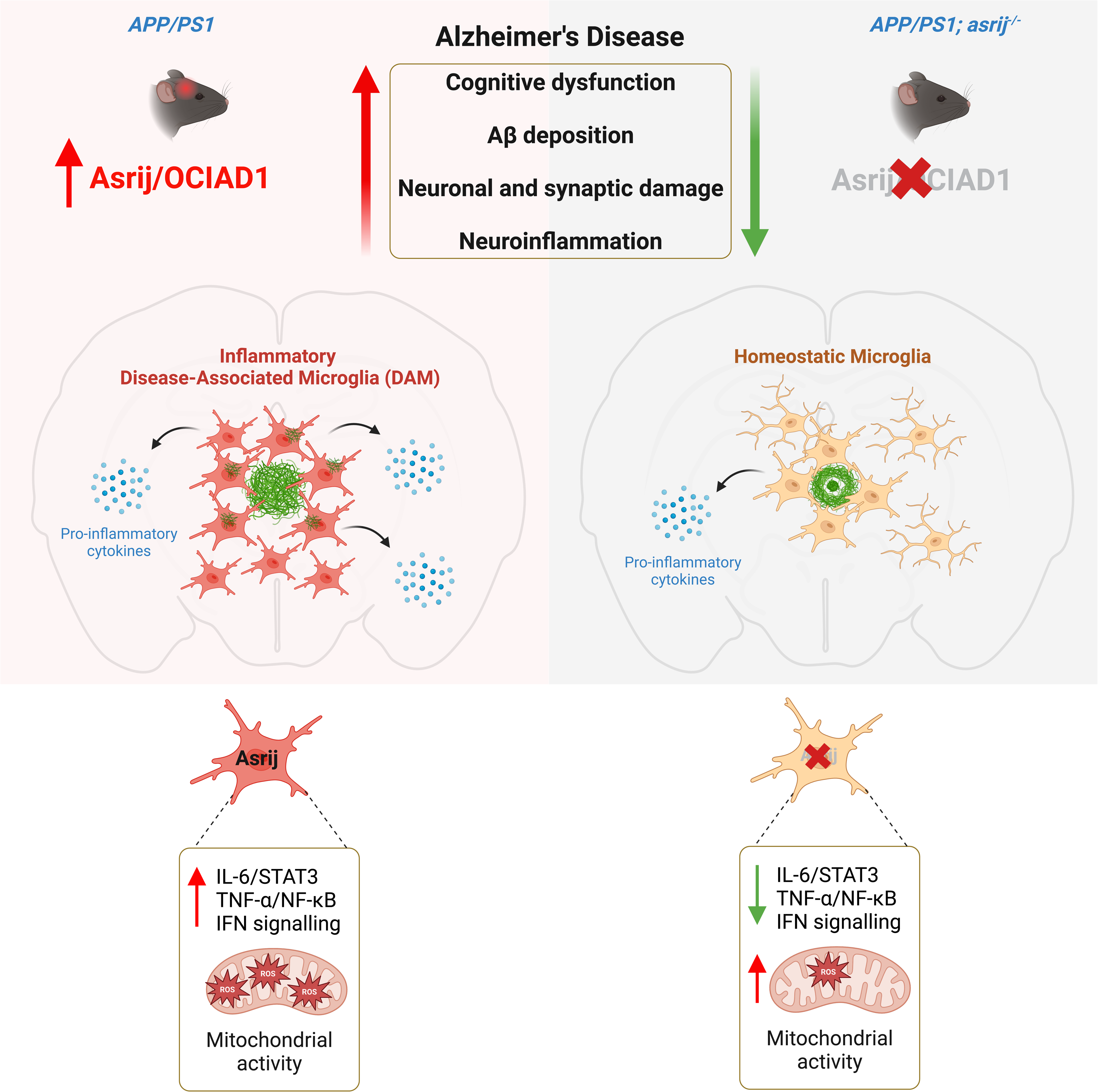

## Background

Alzheimer’s disease (AD) is a chronic, age-related, neurodegenerative disorder and the most common form of dementia. The disease pathology is characterized by the accumulation of extracellular amyloid beta (Aβ) plaques, intracellular neurofibrillary tangles, reactive glia, and neuroinflammation. These pathologies culminate in synaptic loss and neuronal damage, ultimately resulting in learning and memory decline (1). Despite extensive studies on the etiology of AD and its disease pathogenesis, effective treatment strategies are lacking, necessitating the exploration of novel therapeutic targets (2). Genome-wide association studies (GWAS) have identified several genetic loci with variants that alter protein function or expression and are highly implicated in the risk of AD (3). Elucidating the mechanisms by which altered expression of AD-associated factors influence the initiation and progression of disease pathology, may help identify new potential disease-modifying treatments.

Comparative analysis of proteomics and gene expression datasets from AD patients and AD mouse models identified OCIAD1 (Ovarian Carcinoma Immunoreactive Antigen Domain protein 1)/Asrij as a brain vulnerability factor associated with neurodegeneration (4). In AD, Asrij is upregulated via glycogen synthase kinase 3 beta (GSK3β)/β-catenin and p53 signaling and promotes mitochondria-mediated neuronal apoptosis as shown by *in vitro* experiments. AD patients and AD mouse models have increased Asrij levels that correlate with disease severity (4). Moreover, in the AD brain, increased Asrij levels are observed near senile plaques in the afflicted brain regions, namely the hippocampus and cortex, and are associated with dystrophic neurites. This suggests a potential role for Asrij in promoting neurodegeneration in AD. Hence, we tested whether decreasing Asrij levels *in vivo* may mitigate AD phenotypes. Asrij plays critical roles in mammalian blood cell homeostasis (5), stem cell maintenance and differentiation (6,7), and hematopoietic aging (8). Asrij acts as a scaffold protein on endosomes to regulate multiple cellular signaling pathways in *Drosophila* and mice (6,9). It interacts with mitochondrial electron transport chain (ETC) complexes and controls their assembly and activity in human pluripotent stem cells and cell lines (7,10). Moreover, Asrij is essential for mounting immune responses by modulating myeloid cell proliferation and function (5,11). Asrij expression is abundant in the hematopoietic system and brain. In the mouse and human brain, Asrij is expressed predominantly by neurons and microglia with little expression in astrocytes and oligodendrocytes (12) (https://www.proteinatlas.org/ENSG00000109180-OCIAD1/single+cell+type/brain). Despite the early findings linking misexpression of Asrij to AD, whether increased Asrij is a mere consequence of the disease or actively influences disease progression is not known. This motivated us to investigate the *in vivo* role of Asrij in AD pathology.

Here, we show that constitutive depletion of Asrij in the *APP/PS1* mouse model of AD boosts microglial metabolic activity and attenuates pro-inflammatory disease-associated microglia (DAM) responses. This possibly reduces Aβ deposition, decreases neuronal and synaptic loss, curtails neuroinflammation, and improves cognition. Thus, Asrij is a crucial regulator of AD pathology.

## Methods

### Mice

Generation of the *asrij* floxed and knockout (KO) mice has been described previously (5). B6.Cg-Tg(APPswe,PSEN1dE9)85Dbo/MmjaxAPP/PS1 (APP/PS1) mice (Stock #034832-JAX) and C57BL/6J mice (Stock #000664-JAX) were purchased from the Jackson Laboratory. *asrij*^−/−^ mice were crossed to *APP*/*PS1* mice to obtain heterozygous *APP*/*PS1*/*asrij*^+/-^ mice. *APP*/*PS1*/*asrij*^+/−^ mice were further crossed to *asrij*^+/−^ mice to generate the experimental mice, *APP*/*PS1*^−^/*asrij*^+/+^ (denoted as wild type [WT]), *APP*/*PS1*^−^/*asrij*^−/−^ (denoted as KO), *APP*/*PS1*^+^/*asrij*^+/+^ (denoted as AD) and *APP*/*PS1*^+^/*asrij*^−/−^ (denoted as AD/KO). All mice were genotyped using polymerase chain reaction (PCR) before experiments. Mice of both sexes were randomly allocated to experimental groups unless otherwise noted. Mice were maintained at the Jawaharlal Nehru Centre for Advanced Scientific Research (JNCASR) animal facility under ambient temperature (23 ± 1.5°C), 12-h light-dark cycle and with *ad libitum* standard rodent chow diet and water. All mice experiments and protocols were approved by the JNCASR institutional animal ethics committee and were conducted in compliance with the guidelines and regulations.

### Mouse behavioral tests

For all behavioral experiments, 10-month-old male mice were used. Prior to experiments, mice were handled for three consecutive days for acclimatization. Mice were transferred to the behavior test room 30 min before the onset of the test. Open field test (OFT) was used to assess the exploratory and anxiety-like behavior as described previously (13). Briefly, each mouse was introduced into a 50 cm × 50 cm × 20 cm chamber and allowed to move freely for 5 min. 70% ethanol was used to clean the OFT apparatus between consecutive trials. The Morris water maze (MWM) test was used to evaluate learning and memory, as described previously (13). Briefly, a circular water pool of 122 cm diameter and 90 cm depth was filled with water up to 60 cm, and the maze was virtually divided into four quadrants. A 14 cm^2^ platform was placed submerged in one quadrant. Non-toxic white paint was added to the water to keep the tank opaque, and the pool temperature was maintained at 25°C throughout the experiment. During the first five days of training (acquisition), each mouse was subjected to four trials per day with a different starting location for each trial. In each trial, the mouse was placed into the pool facing the wall and allowed 1 min to find the hidden platform. If unable to find the platform, the experimenter would guide the animal towards the platform and let it sit for 20 s before returning to the home cage, and the latency will be recorded as 60 s. On day 7, the escape platform was removed, and the swimming trajectories of the mice during the 60 s trial were recorded. For behavioral tests, mice activity was recorded using an overhead video camera (Sony, #SSC-G118) and analyzed using Smart v3.0 video tracking software (Panlab).

### Immunohistochemistry

Mice were euthanized by cervical dislocation, and brains were swiftly isolated and bisected in ice-cold phosphate-buffered saline (PBS). The left hemisphere underwent fixation in 4% paraformaldehyde (PFA) for 48 h at 4°C. Hippocampi and cortices were micro-dissected from the right hemisphere and flash frozen for storage at −80°C for subsequent protein and RNA extraction. Post fixation, hemibrains were transferred to 30% sucrose for cryoprotection and rehydration for 48 h at 4°C until sunken in the solution. Samples were mounted and frozen in PolyFreeze tissue freezing medium (Sigma-Aldrich, #SHH0026). 30 μm coronal cryosections were obtained using a cryostat (Leica, #CM3050S) and collected on 0.3% gelatin-coated glass slides, and slides were stored at −80°C until further use. For immunofluorescence staining, slides were thawed to room temperature (RT) and washed twice for 5 min with PBS + 0.05% sodium azide, followed by permeabilization with 0.3% Triton X-100 (Sigma-Aldrich, #T8787) in PBS for 1 h at RT. Sections were blocked in 4% fetal bovine serum (FBS) (Gibco, #10270106) for 1 h at RT and then incubated with primary antibodies diluted in blocking solution overnight at 4°C. The following primary antibodies were used: Aβ (D54D2) (Cell Signaling Technology, #8243, 1:300), Aβ (6E10) (BioLegend, #803001, 1:200), LAMP1 (Abcam, #ab24170, 1:200), GFAP (Invitrogen, #130300, 1:200), S100B (Cell Signaling Technology, #90393S, 1:100), IBA1 (Synaptic Systems, #234 009, 1:200), CD68 (Cell Signaling Technology, #97778S, 1:100), Ki67 (Cell Signaling Technology, #9129S, 1:50), CLEC7A (InvivoGen, #mabg-mdect, 1:50) and TMEM119 (Synaptic Systems, #400 002, 1:200). After primary antibody incubation, sections were washed thrice for 10 min with PBS + 0.05% Triton X-100 and then incubated with 1:400 diluted Alexa fluor 488, 568 or 633 conjugated anti-rabbit, -mouse, -rat and -chick secondary antibodies (Invitrogen) as appropriate, diluted in blocking solution for 1 h at RT. Sections were again washed thrice for 10 min with PBS + 0.05% Triton X-100. To visualize plaques, sections were stained with 1% Thioflavin S (ThioS) (Sigma-Aldrich, #T1892) solution in 50% ethanol for 8 min at RT, followed by three 2-min washes with 50% ethanol. After one PBS wash, sections were mounted in ProLong gold antifade mountant (Invitrogen, #P36930) containing 4′,6-diamidino-2-phenylindole (DAPI) (Roche, #10236276001, 1:500) with coverslips.

### Immunoblotting

Frozen hemibrains were thawed on ice and mechanically homogenized using tissue tearor (Biospec Products, #985370) in 500 μL western blot lysis buffer containing 20 mM 4-(2-hydroxyethyl)-1-piperazineethanesulfonic acid (HEPES) (pH = 7.5), 150 mM NaCl, 5 mM MgCl_2_, 5 mM ethylenediaminetetraacetic acid (EDTA) (pH = 8), 0.5% Triton X-100, 10% glycerol, 5 mM dithiothreitol (DTT), 1 mM phenylmethylsulfonyl fluoride (PMSF), 10 mM NaF, 1 mM Na_3_VO_4_ and 1X protease inhibitor cocktail (Sigma-Aldrich, #P8340) on ice. The homogenates were incubated on an end-to-end rotor for 4 h at 4°C for lysis and then spun down at 15,000 rpm for 30 min at 4°C. The supernatant was stored at −80°C, and total protein was estimated using Bradford assay (Bio-Rad, #5000006). For western blotting, equal amounts of protein (40 μg) were denatured in 1X Laemmli buffer containing 5% β-mercaptoethanol at 99°C for 5-10 min. Samples were resolved on 10-12% sodium dodecyl sulfate (SDS) polyacrylamide gel at 150V for 2 h using Mini-PROTEAN tetra cell (Bio-Rad, #1658004). Proteins were transferred on nitrocellulose membrane at 100V for 2 h using the Mini trans-blot cell (Bio-Rad, #1703930). Membranes were blocked with 5% skim milk diluted in PBST (PBS + 0.1% Tween 20) for 1 h at RT on a rocker. Following primary antibodies were diluted in PBST and used at 4°C overnight: APP (Abcam, #ab32136, 1:1000), OCIAD1 (Abcam, #ab91574, 1:1000), GFAP (Cloud-Clone Corp., #PAA068Mu02, 1:1000), Iba1 (Cell Signaling Technology, #17198S, 1:1000), CD68 (Cell Signaling Technology, #97778S, 1:1000), PSD95 (Cell Signaling Technology, #3450S, 1:1000), Synaptophysin (Cell Signaling Technology, #36406S, 1:2000), STAT3 (Cell Signaling Technology, #9139S, 1:1000), Phospho-STAT3 (Tyr705) (Cell Signaling Technology, #9145S), NF-κB p65 (Cell Signaling Technology, #8242S, 1:1000), Phospho-NF-κB p65 (Ser536) (Cell Signaling Technology, #3033S, 1:1000), GAPDH (Sigma-Aldrich, #G9545, 1:4000) and α-tubulin (Sigma-Aldrich, #T8203, 1:2000). Blots were washed thrice for 5 min in PBST and then probed with 1:2000 diluted horseradish peroxidase (HRP)-conjugated anti-rabbit and anti-mouse secondary antibodies (GeNei) for 1 h at RT. After three 5-min washes, blots were developed on X-ray films using Clarity western enhanced chemiluminescence (ECL) reagent (Bio-Rad, #1705061). Band intensities were quantified using Fiji.

### Isolation of microglia from mouse brain

Whole brains isolated from 10-month-old mice were immediately minced into small pieces with a sterile surgical blade in ice-cold Hanks’ balanced salt solution (HBSS) (Thermo Fisher Scientific, #14175095). Tissue chunks were dissociated by mechanical homogenization in 10 mL 1X HBSS containing 15 mM EDTA, 1.5 mM HEPES, and 0.5% glucose by 6-8 gentle strokes in a glass Dounce homogenizer. Homogenates were filtered through 70 μm cell strainers to obtain single-cell suspensions and spun down at 1200 rpm for 10 min at 4°C. Myelin removal was done using Percoll density gradient centrifugation. Briefly, 100% Percoll solution was prepared by mixing absolute Percoll (GE Healthcare, #GE17-0891-02) in 10X Dulbecco’s PBS (DPBS) (Thermo Fischer Scientific, #14080055) at a 9:1 ratio and further diluted to 37% using 1X DPBS. Cell pellets were resuspended in 37% isotonic Percoll followed by centrifugation at 2200 rpm for 15 min at 4°C with minimum acceleration and no brake. The supernatant and myelin layer were aspirated and discarded, and the pellets were washed once with DPBS. Microglia were isolated using Magnetic-activated cell sorting (MACS). Briefly, cells were resuspended in 100 μL MACS buffer (PBS (pH 7.2) + 0.5% bovine serum albumin (BSA) + 2 mM EDTA + 0.05% sodium azide) and incubated with 25 μL CD11b (Microglia) microbeads (Miltenyi Biotec, #130-093-634) on ice for 30 min. Labeled cell suspensions were passed through pre-rinsed LS columns (Miltenyi Biotec, #130-042-401) on a MidiMACS separator magnet (Miltenyi Biotec, #130-042-302), and columns were washed with 5 mL MACS buffer. After removing the columns from the magnet, column-bound cells were plunged with 3 mL MACS buffer and used as purified CD11b^+^ microglia.

### Flow cytometry

Dissociated cells obtained after Percoll density gradient centrifugation were resuspended in 100 μL FACS buffer (PBS (pH 7.2), 1 % BSA + 1 mM EDTA + 0.05% sodium azide) and incubated with CD45-APC-Cy7 (BioLegend, #103116) and CD11b-FITC (BioLegend, #101205) antibodies at 1:100 dilution on ice for 1 h. Cells were washed with FACS buffer at 5000 rpm for 3 min. For analysis of mitochondrial parameters, cells were stained with 25 nM Mitotracker Deep Red (Invitrogen, #M22426) or 50 nM Tetramethylrhodamine, Methyl Ester, Perchlorate (TMRM) (Invitrogen, #T668), or 5 μM MitoSOX Red (Invitrogen, #M36007) diluted in PBS at 37 °C for 30 min. Cells were washed with FACS buffer and analyzed using a BD FACSAria Fusion flow cytometer (BD Bioscience). Debris was excluded using forward and side-scatter pulse area parameters (FSC-A and SSC-A), followed by singlet gating using forward and side-scatter pulse width parameters (FSC-H and SSC-W). Microglia were gated as CD11b^+^ CD45^int^ and median fluorescence intensity (MFI) was quantified. All analysis was carried out using FlowJo v10 software (BD Bioscience).

### *In vitro* Aβ phagocytosis assay

To prepare Aβ oligomers, 1 mg lyophilized Aβ (1–42), HiLyte™ Fluor 555-labeled peptide (AnaSpec, #AS-60480-01) was reconstituted in 1% ammonia solution (NH_4_OH) and then diluted to prepare 100 μM stock solution. 100 μM of monomeric Aβ was incubated for 24 h at 4°C in Ham’s F12 media (Gibco, #11765054) to make a 50 μM stock of oligomeric Aβ as published previously (14,15). For Aβ uptake assay, microglia were incubated with 5 μM oligomeric HiLyte™ Fluor 555 labeled Aβ in Iscove’s modified Dulbecco’s medium (IMDM) (Gibco, #12440053) supplemented with 10% FBS for 1 h at 37 °C. Cells were washed once with PBS and spun down at 5000 rpm for 3 min at 4°C. Following staining with CD11b-APC-Cy7 and CD45-FITC antibodies, samples were analyzed using a BD FACSAria Fusion Flow Cytometer (BD Bioscience).

### Confocal imaging and analysis

Tile scan images of whole brain sections were captured using CQ1 confocal image cytometer (Yokogawa Electric) at 20X magnification. Hippocampus and cortex regions were imaged on a Zeiss laser scanning confocal microscope (LSM) 880 (Zeiss) using 20X and 63X objectives with a *z*-interval of 0.5 μm. To measure plaque density and percent staining area, 20X images from 3 serial sections per mouse were analyzed. Using Fiji, the images were exported to 8-bit format following background subtraction (rolling-ball radius, 50 pixels) and percent staining area was recorded after thresholding. Cell count analysis was performed using the ‘analyze particles’ function in Fiji. For analysis requiring high magnification images, a 63X oil-immersion objective was used with 2X optical zoom and a *z*-interval of 0.5 μm. A minimum of 10-15 individual plaques per brain section were imaged with 3 sections per mouse. To measure microglial Aβ engulfment and plaque volume, 3D surface renderings were created for IBA1, CD68, and ThioS using the ‘surface’ module of Imaris 9.5 software (Bitplane). CD68 and ThioS overlapping surfaces were visualized in a separate channel, and volume was recorded. To assess plaque sphericity, 3D renderings of plaques were analyzed using the built-in ‘sphericity’ function of Imaris, with the value of 1 being equivalent to a perfect sphere. Microglial skeleton analysis was performed using the ‘filament’ module in Imaris. A minimum of 10-15 individual cells per brain section were analyzed with 3 sections per mouse. Maximum intensity projections are shown for all representative images.

### Enzyme-linked immunosorbent assay (ELISA)

For soluble Aβ_1–42_ quantification, supernatants (soluble fractions) collected after homogenization of cortices were used. To extract protein for insoluble Aβ_1–42_ estimation, pellets underwent guanidine extraction. Briefly, brain homogenate pellets were incubated with 5M Guanidine HCl/50 mM Tris (pH = 8.0) in a 1:6 ratio at RT for 3 h. Samples were diluted 1:5 in PBS containing 1X protease inhibitor cocktail and centrifuged at 15,000 rpm for 20 min at 4°C. The supernatant (insoluble fraction) was stored at −80°C until analysis. Protein concentrations of the fractions were determined using Bradford assay. Soluble and insoluble fractions were diluted 1:25 for ELISA. Human Aβ_1–42_ Quantikine ELISA kit (R&D Systems, #DAB142) was used for standard curve generation, and the assay was performed as per the manufacturer’s instructions. To measure cytokine levels in the brain, BD OptEIA mouse IL-6 (BD Biosciences, #550950), BD OptEIA TNF-α (BD Biosciences, #560478) and mouse IL-1β (ABclonal, #RK00006) ELISA kits were used as per manufacturer’s instructions. Absorbance was measured using Varioskan LUX multimode microplate reader (Thermo Fischer Scientific).

### RNA isolation and reverse transcription-quantitative PCR (RT-qPCR)

To isolate RNA, the MACS-sorted CD11b^+^ microglia were resuspended in 500 μL ice-cold TRIzol reagent (Invitrogen, #15596026) and lysed by vortexing. After incubation at RT for 5 min, 100 μL chloroform was added and mixed by gently inverting the tubes. Samples were spun down at 13,000 rpm for 10 min at 4°C. The upper aqueous phase was collected and incubated with an equal volume of 100% isopropanol overnight at −20°C to precipitate RNA. Samples were centrifuged at 13,000 rpm for 20 min at 4°C and the RNA pellet was washed once with 70% ethanol. The pellet was allowed to air dry for 5-10 min at RT and then dissolved in 25 μL ultrapure nuclease-free water (Thermo Fischer Scientific, #10977015). RNA quantity and quality were assessed using a NanoDrop 2000 Spectrophotometer (Thermo Fischer Scientific). cDNA was synthesized using 200-400 ng total RNA in SuperScript III first-strand synthesis kit (Invitrogen, #18080051) following the manufacturer’s instructions. RT-qPCR was set up using SensiFAST SYBR No-ROX kit (Bioline, #BIO-98020) in CFX384 real-time PCR system (Bio-Rad) as per the manufacturer’s protocol. Following primers were used: *asrij* forward 5’-GGAGATCTCGAGATGAATGGGAGGGCTGATTTT-3’ and reverse 5’-GTGGTGGCGGCCGCTCACTCATCCCAAGTATCTCC −3’; *β-actin*forward 5’-GTGACGTTGACATCCGT-3’ and reverse 5’-GCCGGACTCATCGTACTCC-3’. All reactions were set up in duplicates, and *β-actin* was used as the loading control.

### RNA sequencing and analysis

RNA isolated from MACS-sorted microglia by the TRIzol method was quantified using Qubit high-sensitivity assay kit (Invitrogen, #Q32852) on Qubit 4 fluorimeter (Invitrogen). Samples were run on the tapeStation system (Agilent) utilizing high sensitivity RNA ScreenTape analysis (Agilent, #5067-5579) for RNA quality control (QC). Sample QC, library preparation, and RNA sequencing were done by the next generation genomics facility at the Bangalore Life Science Cluster (BLiSC), India. Briefly, ribosomal RNA (rRNA) depletion was performed using NEBNext rRNA depletion kit v2 (Human/Mouse/Rat) with RNA sample purification beads (NEB, #E7405X). cDNA libraries were constructed utilizing the NEBNext Ultra II Directional RNA Library Prep (stranded) kit with sample purification beads (NEB, #E7765L). cDNA libraries were sequenced on the NovaSeq 6000 platform (Illumina) with 2×100bp sequencing read length and ∼100 million read depth per sample. Sequenced reads in FASTQ files were trimmed and filtered to remove adaptor sequences and low-quality bases, respectively, using the fastp tool. Data quality was checked using the FastQC tool. Reads were aligned to the indexed UCSC mm10 mouse reference genome using HISAT 2. SAM files were converted and sorted/indexed by coordinate to BAM files using Samtools. FeatureCounts program of the Subread R package was used to calculate and summarize per-gene read counts as expression values. DESeq2 (v1.30.0) was utilized for differential gene expression (DEG) analysis. Principal component analysis (PCA) plots were generated using the plotPCA command in DEseq2. DEGs (p < 0.05) with log_2_fold change > 0.5 were considered upregulated, and log_2_fold change < −0.5 were considered downregulated. Volcano plots were generated using the ggplot2 package. Heatmaps were visualized using the pheatmap R package. For analysis and visualization of gene ontology (GO) enrichment terms, the clusterProfiler R package was used at default settings. Pathway enrichment was conducted using the Enrichr web tool and MSigDB collection.

### Statistical analysis

Each data point in the graphs represents an individual mouse. All data are presented as mean ± standard error of mean (SEM). Statistical significance between the two groups was determined using a two-tailed unpaired Student’s *t*-test. Comparisons between more than two groups were made using one-way, two-way, or repeated measures analysis of variance (ANOVA) with Bonferroni’s or Tukey’s multiple comparisons post-hoc test. All statistical analyses and graph plotting were performed in Prism 10 software (GraphPad). *P* values less than 0.05 were considered significant. ns - not significant, \**P* < 0.05, \*\**P* < 0.01, and \*\*\**P* < 0.001.

## Results

### Asrij depletion alleviates cognitive deficits and anxiety-like behavior in *APP/PS1* mice

To investigate how Asrij impacts Aβ pathology and Alzheimer’s disease (AD)-related behavior, we crossed *asrij*^−/−^ (knockout [KO]) mice with *APP/PS1* mice, a well-studied mouse model of AD, to generate *APP/PS1*^+^ *asrij*^−/−^ (referred to as AD/KO) mice. We used *APP/PS1*^−^ *asrij*^+/+^ (referred to as wild type [WT]) mice, *APP/PS1*^−^ *asrij*^−/−^ (referred to as KO) mice, and *APP/PS1*^+^ *asrij*^+/+^ (referred to as AD) mice as controls (Fig. S1A-B). Immunoblotting analysis confirmed that Asrij levels were increased in AD mice and were undetectable in AD/KO mice (Fig. S1C).

*APP/PS1* mice exhibit learning and memory deficits from 9 months of age (16). To evaluate the effect of Asrij depletion on cognitive performance, we performed the Morris water maze (MWM) test using 10-month-old mice (Fig. 1A). Over the first 5 days of the training phase, WT and KO mice showed gradually decreasing latency to find the submerged platform, which indicates intact spatial learning. AD mice took a substantially longer time searching for the hidden platform. Interestingly, AD/KO mice showed significantly reduced latency to find the platform as compared to AD mice, suggesting improved spatial learning abilities (Fig. 1B). The swimming path used by the mice to search for the hidden platform during the memory recall session on day 7 also reflects the extent of cognitive dysfunction. As seen from mouse path traces, AD mice adopted a random and non-directed navigation path, but AD/KO mice performed a focused and self-orienting search for the platform. (Fig. 1C). Moreover, AD/KO mice showed reduced latency to the platform, increased crossing number, and spent significantly more time in the target quadrant, demonstrating restored spatial memory (Fig. 1D-F). Importantly, differences in performance in the MWM test did not arise due to defects in locomotor activity as comparable swimming speeds were noted for all the genotypes (Fig. 1G). We did not find any appreciable difference in MWM test parameters between WT and KO mice suggesting that Asrij depletion does not affect learning and memory, in the absence of Aβ pathology. Thus, Asrij deficiency alleviates learning and memory defects in *APP/PS1* mice.

**Figure 1.**
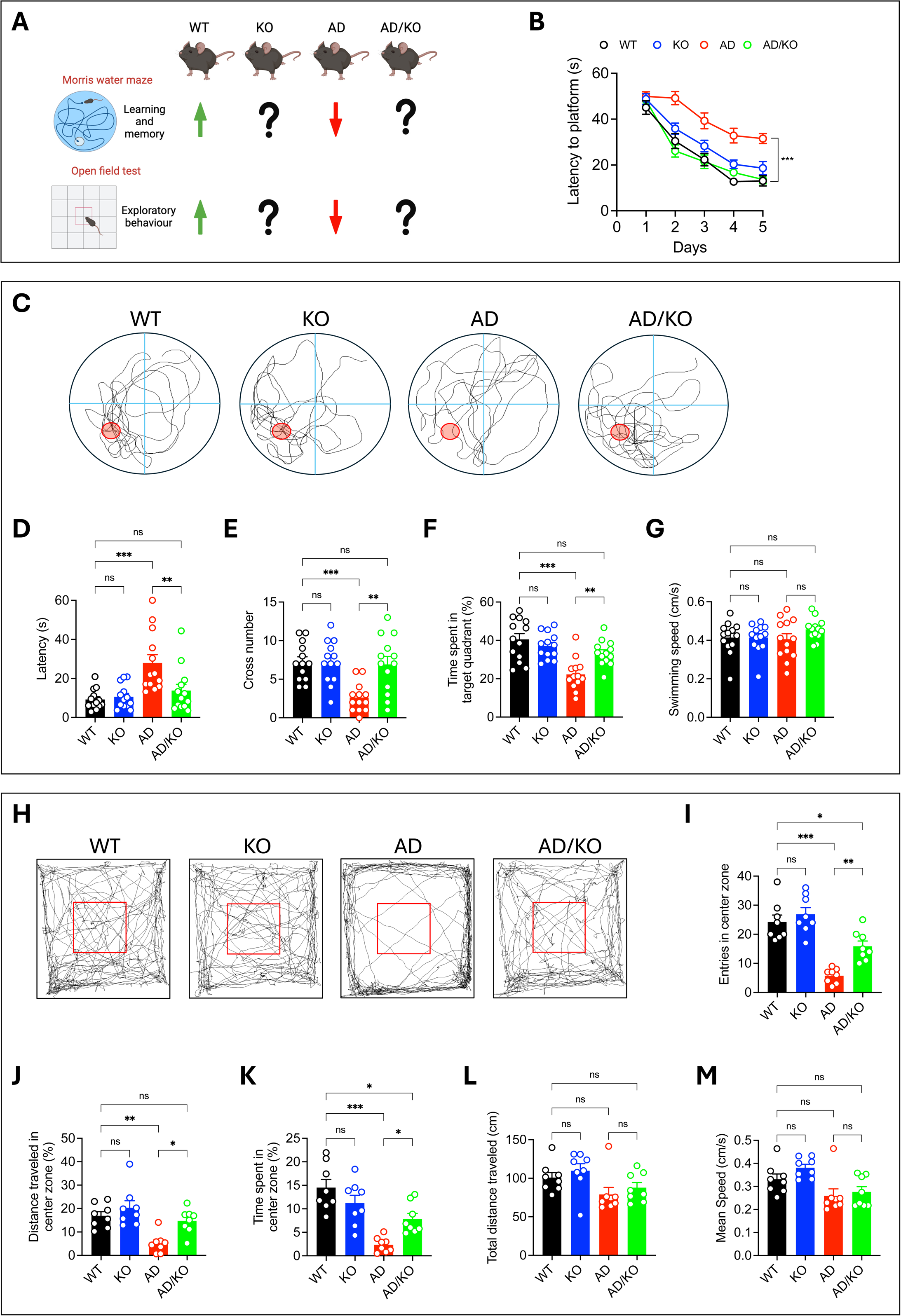
Asrij depletion alleviates cognitive deficits and anxiety-like behavior in *APP/PS1* mice. (A) Schematic showing behavioral tests used to assess cognitive performance of mice. (B) Graph shows quantification of latency to platform across five days of invisible platform training (acquisition) in Morris water maze (MWM) test (n = 13) (C) Representative images of mouse tracks during MWM probe trial. The red circle indicates the location of the platform during training. Graphs show quantification of (D) latency, (E) cross number, (F) the percentage of time spent in the target quadrant, and (G) swimming speed (cm/s) during the MWM probe trial (n = 13). (H) Representative traces of mouse path in open field test (OFT). Graphs show quantification of (I) entries in the center zone, (J) the percentage of distance traveled in the center zone, (K) the percentage of time spent in the center zone, (L) total distance traveled, and (M) mean speed (cm/s) in OFT (n = 8). Statistical significance between experimental groups was calculated by repeated-measures two-way ANOVA with Bonferroni’s post hoc test (B) and one-way ANOVA with Bonferroni’s post hoc test (D-G and I-M). Error bars denote the mean ± SEM. ns-non-significant, \**P* < 0.05, \*\**P* < 0.01, and \*\*\**P* < 0.001.

Aβ pathology is also known to reduce exploratory behavior and spur anxiety-like behavior in *APP/PS1* mice, wherein they tend to spend more time in the periphery of an open field, referred to as thigmotaxis, a measure of anxiety (16). Therefore, we evaluated the effect of Asrij depletion on these behaviors using the open field test (OFT). During the 5-minute free exploration period, AD mice had reduced entries in the center, spent less time, and covered less distance in the center zone of an open field compared to WT mice, suggesting elevated anxiety. Interestingly, AD/KO mice showed increased center entries and increased time spent and distance covered in the center (Fig. 1H-K). Asrij depletion in the absence of Aβ pathology did not affect performance in the OFT as there was a comparable level of exploratory behavior between WT and KO mice. Further, no differences were recorded in the total distance traveled and speed of mice of all four genotypes, suggesting that results obtained in OFT were not due to altered locomotor activity (Fig. 1L-M). Thus, Asrij depletion improves exploratory behavior and limits anxiety-like phenotypes in *APP/PS1* mice. Taken together, our results suggest that an increase in Asrij levels in AD potentially precipitates neurobehavioral and cognitive dysfunction in AD.

### Asrij depletion reduces Aβ plaque deposition and associated pathology in *APP/PS1* mice

In the *APP/PS1* mice, Aβ plaque deposition and neuroinflammation begins at around 4 months and progresses gradually, resulting in a full-blown disease by 10 months of age (17). To determine whether Asrij influences the accumulation of Aβ plaques, we first stained brain cryosections with Thioflavin S (ThioS), which marks the dense core compact Aβ plaques. We did not observe a significant difference in plaque load between 5-month-old AD and AD/KO mice (Fig. S2A). However, 10-month-old AD/KO mice exhibited a remarkable reduction in Aβ plaques in the cortex and hippocampus compared to AD mice (Fig. 2A). In parallel, we also assessed the accumulation of total Aβ in the brain using the D54D2 antibody, which recognizes the total Aβ peptide burden including various isoforms such as the most abundant Aβ_40_ (relatively inert) and Aβ_42_ (highly neurotoxic) species. Immunostaining analysis revealed significantly reduced total Aβ levels in the cortex and hippocampus of AD/KO mice (Fig. 2B). This indicates that Asrij depletion limits Aβ plaque accumulation in *APP/PS1* mice.

**Figure 2.**
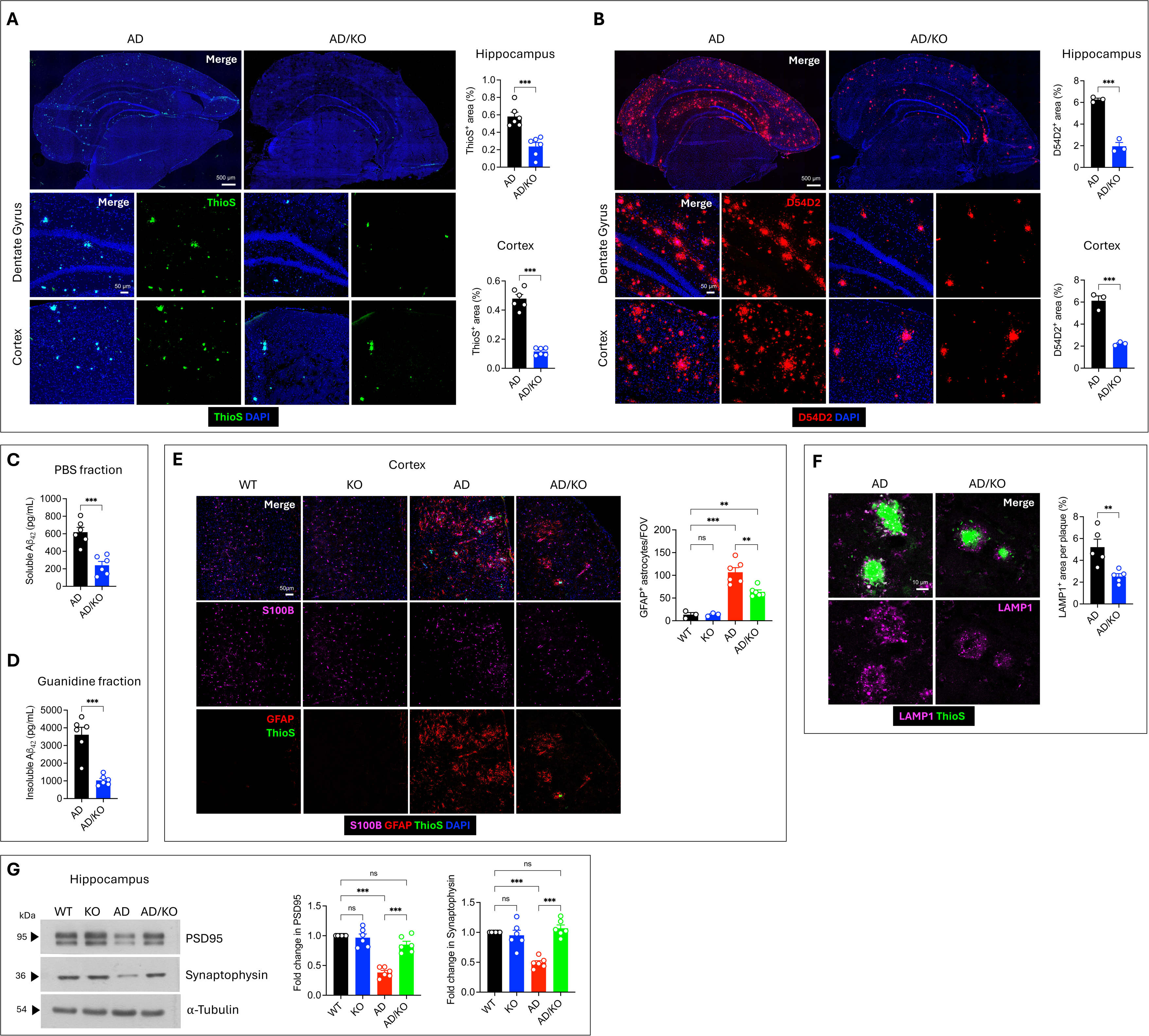
Asrij depletion reduces Aβ plaque deposition and associated pathology in *APP/PS1* mice. (A) Representative confocal images of brain sections showing Thioflavin S (ThioS) staining (green) for plaques. DAPI (blue) marks the nuclei. Graphs show quantification of percent area covered by ThioS^+^ staining in hippocampus and cortex (n = 6). (B) Representative confocal images of brain sections showing Aβ staining (red) using D54D2 antibody. DAPI (blue) marks the nuclei. Graphs show quantification of percent area covered by Aβ staining in hippocampus and cortex (n = 3). (C, D) Graphs show quantification of Aβ_42_ levels (pg/mL) in the soluble (PBS) and insoluble (guanidine) brain fractions by ELISA (n = 6). (E) Representative confocal images showing LAMP1 staining (magenta) for dystrophic neurites around ThioS^+^ Aβ plaques (green). Graph shows quantification of LAMP1^+^ area per plaque (n = 5). (F) Immunoblot analysis of PSD95 and Synaptophysin levels in the hippocampus. α-tubulin is loading control. Graphs show fold change in PSD95 and Synaptophysin levels normalized to α-tubulin (n = 6). (G) Representative confocal images showing S100B (magenta) and GFAP (red) staining for astrocytes in the cortex. ThioS (green) and DAPI (blue) mark plaques and nuclei, respectively. Graph shows quantification of GFAP^+^ astrocytes per field of view (FOV) of the cortex (n = 3 for WT and KO, n = 6 for AD and AD/KO). Statistical significance between experimental groups was calculated by unpaired two-tailed Student’s t-test (A-E) and one-way ANOVA with Tukey’s post hoc test (F-G). Error bars denote mean ± SEM. ns-non-significant, \**P* < 0.05, \*\**P* < 0.01, and \*\*\**P* < 0.001.

Soluble Aβ oligomers, which are highly neurotoxic, aggregate to form insoluble fibrils and, subsequently, compact plaques (18). To evaluate whether the absence of Asrij alters the soluble and insoluble forms of Aβ in AD mice, we quantified the concentration of Aβ_42_ species in the soluble (phosphate buffered saline (PBS)-extracted) and insoluble (guanidine hydrochloride-extracted) brain fractions using the enzyme-linked immunosorbent assay (ELISA). In alignment with the reduced Aβ staining observed in imaging studies, we detected substantially reduced amounts of Aβ_42_ in soluble and insoluble brain fractions (Fig. 2C-D). Notably, the levels of the full-length amyloid precursor protein (APP) and its cleaved carboxy-terminal fragments (CTF), CTF-α and CTF-β were unchanged, suggesting that reduction in plaques upon Asrij depletion does not result from altered APP processing (Fig. S2B). Altogether, these studies strongly support the idea that Asrij promotes Aβ plaque accumulation and that its absence reduces the overall Aβ burden in *APP/PS1* mice.

One of the major pathological features of AD is the presence of hypertrophic reactive glial fibrillary acidic protein (GFAP) positive astrocytes around Aβ plaques, that are known to propagate neurotoxicity (19). To visualize reactive astrogliosis, we performed immunofluorescence staining for GFAP, an astrocyte marker whose levels increase upon activation, and S100 calcium-binding protein B (S100B), a pan-astrocyte marker. Note that cortical astrocytes do not express GFAP under homeostatic conditions, but in conditions of disease or injury, GFAP immunoreactivity in the cortex represents astrocyte activation (19). Indeed, we found a marked increase in the number of GFAP^+^ astrocytes in the cortex of AD mice. AD/KO mice showed reduced GFAP^+^ astrocytes in the cortex as compared to AD mice (Fig. 2E). Consistently, immunoblotting analyses showed reduced GFAP protein levels (Fig. S2C). Thus, Asrij depletion attenuates Aβ associated astrocyte activation, in line with the reduced neuropathology.

Extracellular Aβ deposits are surrounded by swollen dystrophic axons and dendrites of neurons, called dystrophic neurites, that are marked by abnormal accumulation of lysosomal-associated membrane protein 1 (LAMP1) vesicles (20). Given the improved behavioral performance and reduced Aβ deposition in AD/KO mice, we next quantified LAMP1 immunoreactivity. We found that plaques in AD/KO mice were covered by reduced LAMP1 staining, suggesting attenuated neuronal dystrophy (Fig. 2F). Additionally, the absence of Asrij restored levels of post-synaptic marker, post synaptic density 95 (PSD95) and pre-synaptic marker, Synaptophysin, indicating synaptic protection (Fig. 2G). Collectively, these results show that Asrij depletion prevents Aβ induced neuronal and synaptic damage.

### Reduced microgliosis and microglial Aβ phagocytosis in Asrij deficient *APP/PS1* mice

Microglia are brain-resident innate immune cells that play beneficial and detrimental roles in a context-specific manner in AD (21). In response to Aβ, microglia undergo activation, wherein they are recruited to the plaques, undergo local proliferation, and form a physical barrier to protect neurons from Aβ-mediated toxicity (22). Microglial Aβ phagocytosis contributes to Aβ clearance (23) but also promotes the formation and deposition of Aβ plaques (24–26). Interestingly, publicly available datasets revealed marginally increased Asrij expression in AD microglia (Fig. S3A-B). We also performed reverse transcriptase-quantitative polymerase chain reaction (RT-qPCR) and immunoblotting analyses for Asrij on magnetic-activated cell sorting (MACS)-sorted CD11b^+^ mouse microglia. Consistently, AD microglia had increased transcript and protein levels of Asrij, compared to WT microglia (Fig. S3C-D). This hints that Asrij may be necessary in regulating microglial responses to Aβ pathology. Additionally, Asrij controls myeloid cell function and immune responses in *Drosophila* and mice (5,11). Increased Asrij levels in AD microglia, combined with reduced Aβ pathology, led us to hypothesize that Asrij may alter microglial activity in AD mice.

To evaluate microgliosis and phagocytic activation, we performed immunofluorescence staining for a microglial marker, ionized calcium-binding adapter molecule 1 (IBA1), and a phagolysosomal marker, cluster of differentiation 68 (CD68). AD/KO mice had fewer microglia in the hippocampus and cortex than AD mice. Further, CD68 coverage was significantly reduced in AD/KO mice, confirming lower microglial phagocytic activation (Fig. 3A-B). In agreement with our imaging results, immunoblotting analysis showed that IBA1 and CD68 levels were reduced in AD/KO mice (Fig. S4A). Low levels of microgliosis and phagocytic activation upon Asrij depletion appears to be specific to Aβ pathology as there was no significant difference in microglial numbers between WT and KO mice. We also observed reduced microglial clustering to Aβ plaques as seen by a two-fold reduction in the number of plaque-associated microglia in AD/KO as compared to AD mice (Fig. 3C). Proliferation is known to contribute to localized increase in microglial numbers around plaques (22). Ki67 staining for assessing proliferation showed that AD/KO mice had fewer Ki67^+^ IBA1^+^ plaque-proximal microglia than AD mice, indicating attenuated microglial proliferation (Fig. 3D). Thus, Asrij is required for microglial recruitment, proliferation and association with Aβ plaques.

**Figure 3.**
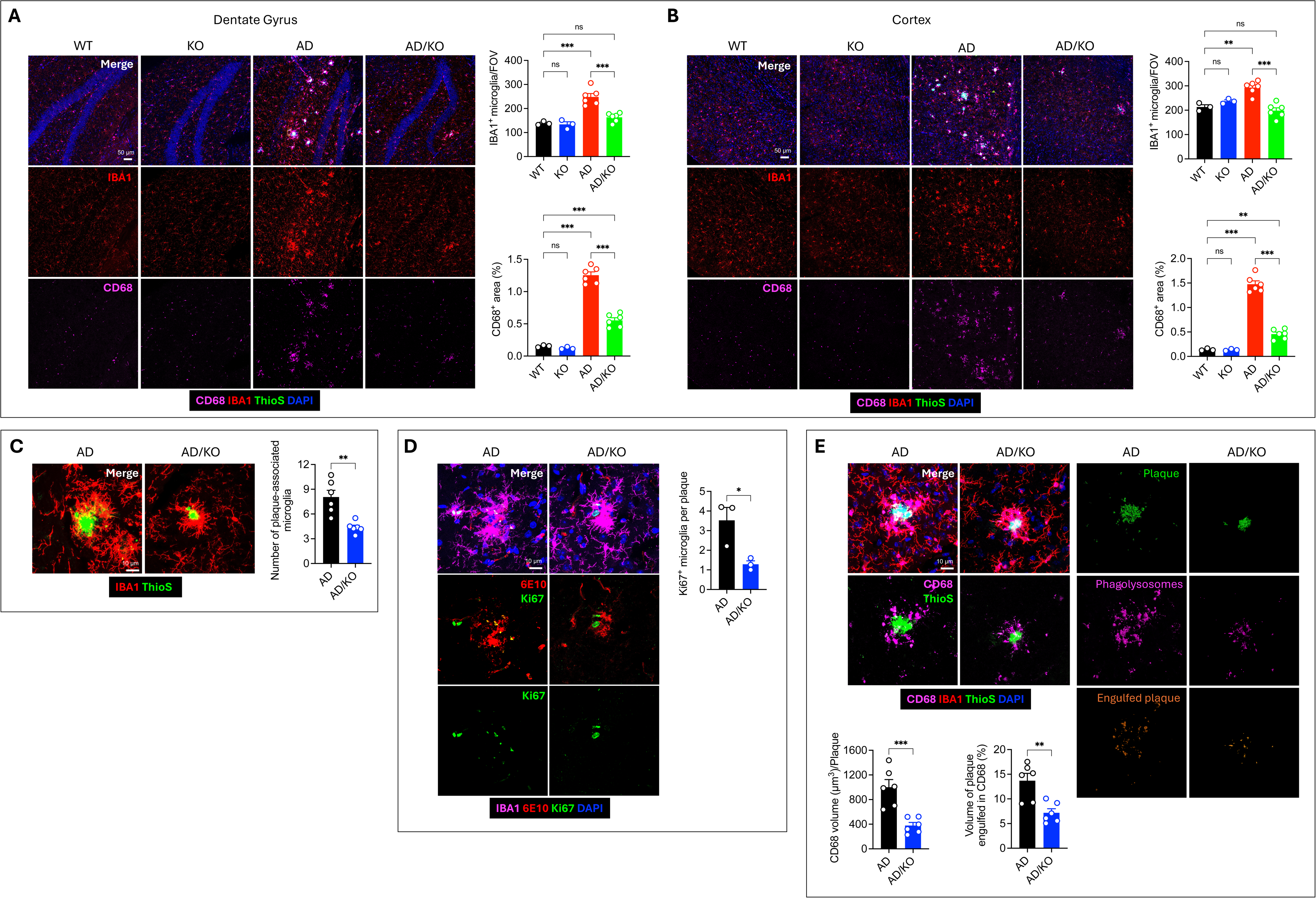
Reduced microgliosis and microglial Aβ phagocytosis in Asrij deficient *APP/PS1* mice. (A) Representative confocal images showing IBA1^+^ microglia (red), CD68^+^ phagolysosomes (magenta) and ThioS^+^ Aβ plaques (green) in the hippocampus. DAPI (blue) marks the nuclei. Graphs show quantification of microglial numbers per field of view (FOV) and percent area covered by CD68 staining in the hippocampus (n = 3 for WT and KO, n = 6 for AD and AD/KO). (B) Representative confocal images showing IBA1^+^ microglia (red), CD68^+^ phagolysosomes (magenta), and ThioS^+^ Aβ plaques (green) in the cortex. DAPI (blue) marks the nuclei. Graphs show quantification of microglial numbers per FOV and percent area covered by CD68 staining in the cortex (n = 3 for WT and KO, n = 6 for AD and AD/KO). (C) Representative confocal images showing IBA1^+^ microglia (red) around ThioS^+^ Aβ plaque (green). Graph shows quantification of the number of microglia within a 15 μm radius surrounding Aβ plaque (n = 6). (D) Representative confocal images showing Ki67^+^ (green) proliferating IBA1^+^ microglia (magenta) associated with 6E10^+^ Aβ plaques (red). DAPI (blue) marks the nuclei (n = 3). Graph shows quantification of the number of Ki67^+^ microglia per plaque. (E) Representative confocal images and Imaris-based 3D rendering of ThioS^+^ Aβ plaques (green), IBA1^+^ microglia (red), and CD68^+^ phagolysosomes (magenta). Aβ volume colocalized within the CD68 volume is shown as a separate channel (orange). Graphs show quantification of CD68 staining volume per plaque and the percentage of plaque volume engulfed in CD68 staining (n = 6). Statistical significance between experimental groups was calculated by one-way ANOVA with Tukey’s post hoc test (A-B) and unpaired two-tailed Student’s t-test (C-E). Error bars denote mean ± SEM. ns-non-significant, \**P* < 0.05, \*\**P* < 0.01, and \*\*\**P* < 0.001.

Microglial Aβ phagocytosis shapes the extent of plaque accumulation and is a critical response to AD pathology (27). To evaluate Aβ engulfment by microglia, we utilized Imaris-based 3D rendering of ThioS, IBA1, and CD68 staining. Asrij depleted microglia showed an three-fold reduction in the volume of ThioS^+^ plaques internalized within CD68^+^ phagosomes. Also, plaque-associated CD68 staining was lower in AD/KO mice indicating reduced microglial Aβ engulfment (Fig. 3E). As a secondary approach to assess microglial phagocytosis, we performed flow cytometry-based *in vitro* oligomeric Aβ uptake assay. The frequency of CD11b^+^ CD45^int^ microglia in AD/KO mice was lower than in AD mice, confirming reduced microgliosis observed in imaging studies (Fig. S4B-C). Also, levels of CD45 and CD11b, which are generic markers of activation, were significantly diminished in AD/KO mice (Fig. S4C). We did not observe any difference in microglial frequency and activation between WT and KO mice. Thus, Asrij depletion limits microgliosis and microglial activation in the context of Aβ pathology. When assessing Aβ uptake, approximately 40% of CD11b^+^ CD45^int^ microglia in WT and KO mice were positive for HiLyte Fluor 555 labeled oligomeric Aβ_42_, reflecting unchanged basal phagocytosis activity. As expected, AD microglia showed increased Aβ_42_ uptake while AD/KO microglia exhibited dampened Aβ phagocytosis with about 38% microglia ingesting labeled oligomers (Fig. S4D). These results provide evidence that Asrij critically contributes to the uptake and phagocytosis of Aβ plaques by microglia, in the AD mouse model.

Microglial barrier function mediates the physical compaction of Aβ plaques (20,28). Since we observed reduced Aβ footprint in AD/KO mice, we postulated that Asrij deficient microglia may promote Aβ compaction. To this end, we evaluated plaque volume and plaque sphericity, a readout of microglial barrier function (29). Spherical compact plaques are known to be less neurotoxic than the irregularly shaped, loosely organized fibrillar Aβ plaques (20). We found that plaques in AD/KO mice exhibited more sphericity than those in AD mice, indicating effective compaction (Fig. S5). Additionally, we performed co-immunostaining analysis using 6E10 antibody (that detects filamentous/fibrillar Aβ) and ThioS (that marks inert compact amyloid). Expectedly, Aβ deposits in AD/KO mice had less labeling of 6E10 and more labeling of ThioS, as reflected in the diffuseness index (Area_ThioS_/Area_6E10_) (Fig. S5). These findings support that the absence of Asrij favors the compactness of amyloid plaques and enhances microglial plaque consolidation in *APP/PS1* mice.

### Asrij deficiency upregulates metabolic genes and increases mitochondrial activity in AD microglia

To gain a comprehensive and unbiased assessment of molecular mechanisms by which Asrij impacts microglial homeostasis in AD, we performed bulk RNA sequencing (RNA-seq) on immunomagnetic bead-sorted CD11b^+^ microglia, isolated from the brains of AD and AD/KO mice (Fig 4A). We found 1863 upregulated genes (log_2_ fold change > 0.5) and 623 downregulated genes (log_2_ fold change < −0.5) in AD/KO microglia compared to AD microglia (Fig. 4B and Fig. S6A). Gene ontology (GO) analysis of differentially expressed genes (DEGs) revealed upregulation of biological processes such as mitochondrial ATP synthesis, NADH dehydrogenase complex assembly, protein translation, and phosphatidylcholine metabolism. On the other hand, downregulated biological processes were related to the regulation of innate immune responses, defense response to pathogens, and mitotic cell cycle (Fig. S6B). Consistently, upregulated genes were enriched for GO terms related to mitochondrial respiratory chain complexes, ribosomal subunits, and protein folding while downregulated genes were associated with immune receptor activity, peptidase activity, mitotic spindle and cell cycle (Fig. S6C-D). Pathway enrichment analysis showed activation of metabolic pathways such as oxidative phosphorylation, citric acid cycle, glycolysis, and cholesterol homeostasis, among others (Fig. 4C-D). Altogether, these results indicate that Asrij depletion predominantly results in increased expression of mitochondrial activity genes with a concomitant reduction in innate immunity and proliferation genes in microglia, suggesting a protective response to Aβ pathology.

**Figure 4.**
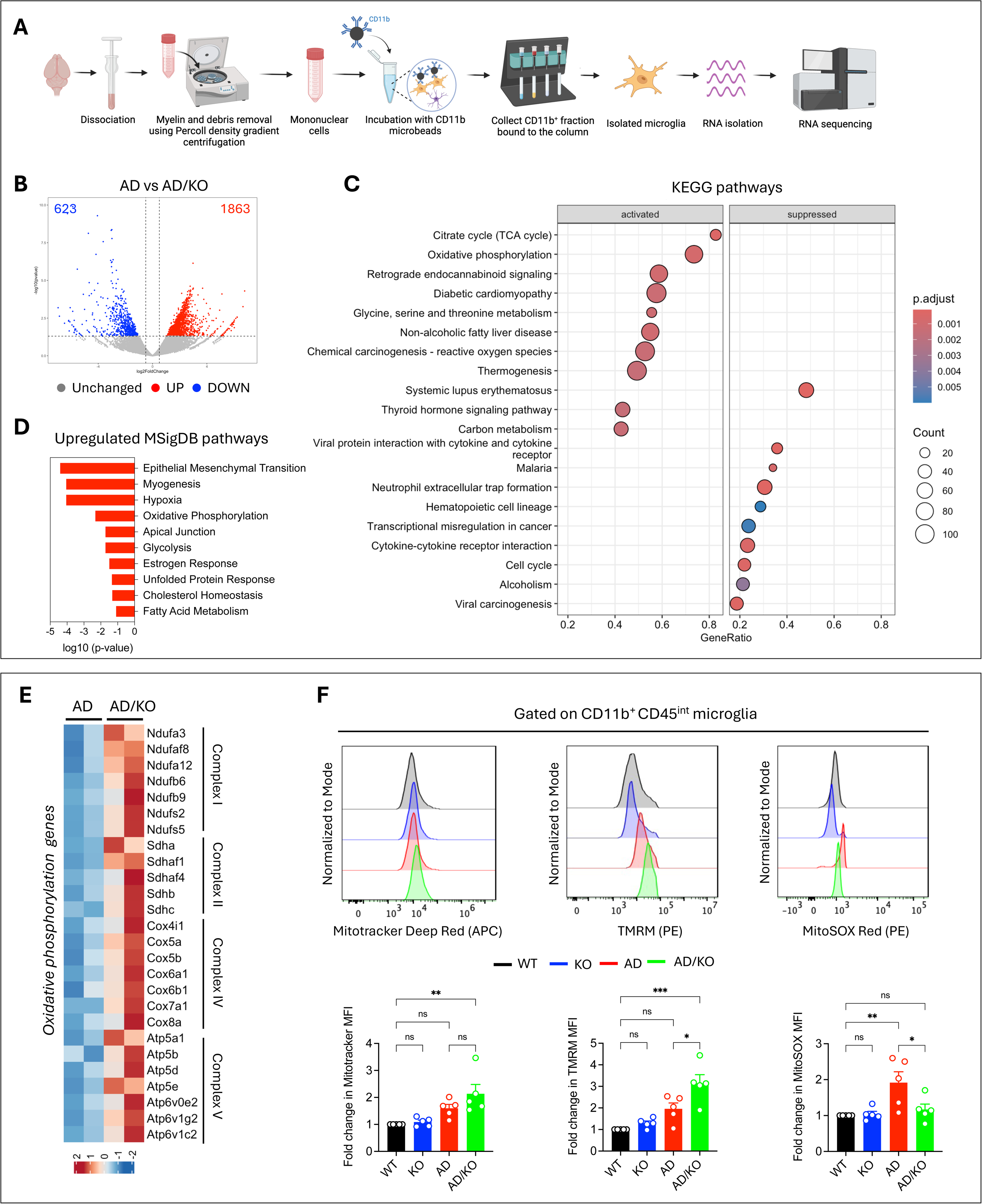
Asrij deficiency upregulates metabolic genes and increases mitochondrial activity in AD microglia. (A) Schematic representing the experimental workflow of microglia isolation for RNA-seq analysis (n = 2 independent experiments). (B) Volcano plot depicting differentially expressed genes (DEGs). Points in red and blue are significantly (*P* < 0.05) upregulated (log_2_FC > 0.5) and downregulated (log_2_FC < −0.5) genes, respectively. (C) KEGG term enrichment dot plot shows the top 8 significantly activated and repressed pathways (adjusted *P* < 0.05) using clusterProfiler R package. (D) Bar graph shows the top 10 significantly upregulated MSigDB hallmark pathways (*P* < 0.05). (E) Heatmap of significantly upregulated (*P* < 0.05, log_2_FC > 0.5) genes involved in oxidative phosphorylation. (F) Representative histograms show flow cytometry analysis of Mitotracker Deep Red, TMRM, and MitoSOX Red staining in CD11b^+^ CD45^int^ microglia. Graphs show quantification of fold change in Median Fluorescence Intensity (MFI) of the indicated mitochondrial dyes (n = 5). Statistical significance between experimental groups was calculated by two-sided Wald test (B), hypergeometric test (C), Fischer exact test (D), and one-way ANOVA with Tukey’s post-hoc test (F). Error bars denote mean ± SEM. ns-non-significant, \**P* < 0.05, \*\**P* < 0.01, and \*\*\**P* < 0.001.

Activated microglia must maintain a high energy metabolism coupled with anabolic processes to counter Aβ pathology (30, 31, 32). Asrij interacts with electron transport chain (ETC) components and regulates Complex I activity and Complex III assembly (7,10). Hence, we chose to investigate changes in mitochondrial activity in greater detail. Firstly, we analyzed the expression of genes that encode mitochondrial ETC complex proteins. AD/KO microglia exhibited upregulation of genes related to subunits of NADH dehydrogenase complex, succinate dehydrogenase, cytochrome c oxidase complex, and ATP synthase (Fig. 4E). Next, we performed flow cytometry to evaluate mitochondrial activity of CD11b^+^ CD45^int^ microglia using mitochondria specific dyes. AD/KO microglia had equivalent staining for Mitotracker Deep Red as compared to AD microglia, suggesting unchanged mitochondrial mass. Interestingly, we observed increased Tetramethylrhodamine, Methyl Ester, Perchlorate (TMRM) intensity in AD/KO microglia indicating increased mitochondrial membrane potential, an indicator of mitochondrial activity. Further, MitoSOX Red staining revealed reduced mitochondrial reactive oxygen species (ROS) in AD/KO microglia (Fig. 4F). Noticeably, lack of Asrij did not have any overt effect on the mitochondrial parameters in the absence of Aβ, suggesting that this could be an AD-specific response.

Upregulation of oxidative phosphorylation-related genes and mitochondrial activation is one of the hallmarks of mammalian target of rapamycin (mTOR) driven increase in microglial metabolism (31,33). Asrij negatively controls Akt activation in bone marrow cells (5). Hence, we assessed the activation status of Akt-mTOR signaling. Immunoblotting analyses on magnetic bead-sorted CD11b^+^ cells demonstrated increased phosphorylation of Akt and mTOR in AD/KO microglia in comparison to AD microglia. No changes were observed in the phosphorylated Akt and mTOR levels between WT and KO mice (Figure S6E). Collectively, these data strongly indicate that Asrij negatively controls mitochondrial activity and the Akt-mTOR signaling axis. Thus, depletion of Asrij possibly promotes microglial metabolic fitness, resulting in a neuroprotective microglial response in AD.

### Asrij promotes inflammatory disease-associated microglia (DAM) activation state

Upon sensing Aβ and inflammatory cues, microglia undergo dramatic morphological changes that reflect their activation status. Homeostatic/resting microglia typically exhibit a small soma with highly ramified processes. Microglial activation is accompanied by a shift to larger soma size and retraction of processes, resulting in the acquisition of an amoeboid morphology (23). This morphological transition is pivotal for the subsequent microglial immune responses, such as phagocytosis and cytokine production that influence Aβ pathology (20,34). Since Asrij depletion reduced microglial activation, we analyzed microglial morphology. AD/KO microglia harbored fewer branches and had reduced total branch length, highlighting reduced morphological complexity compared to AD microglia. In contrast, WT and KO microglia did not exhibit any morphological changes (Fig. 5A). This illustrates that the effects of Asrij depletion on microglial morphology largely manifest in the presence of Aβ pathology. Thus, Asrij is pivotal for microglia to attain a morphologically activated state in AD.

**Figure 5.**
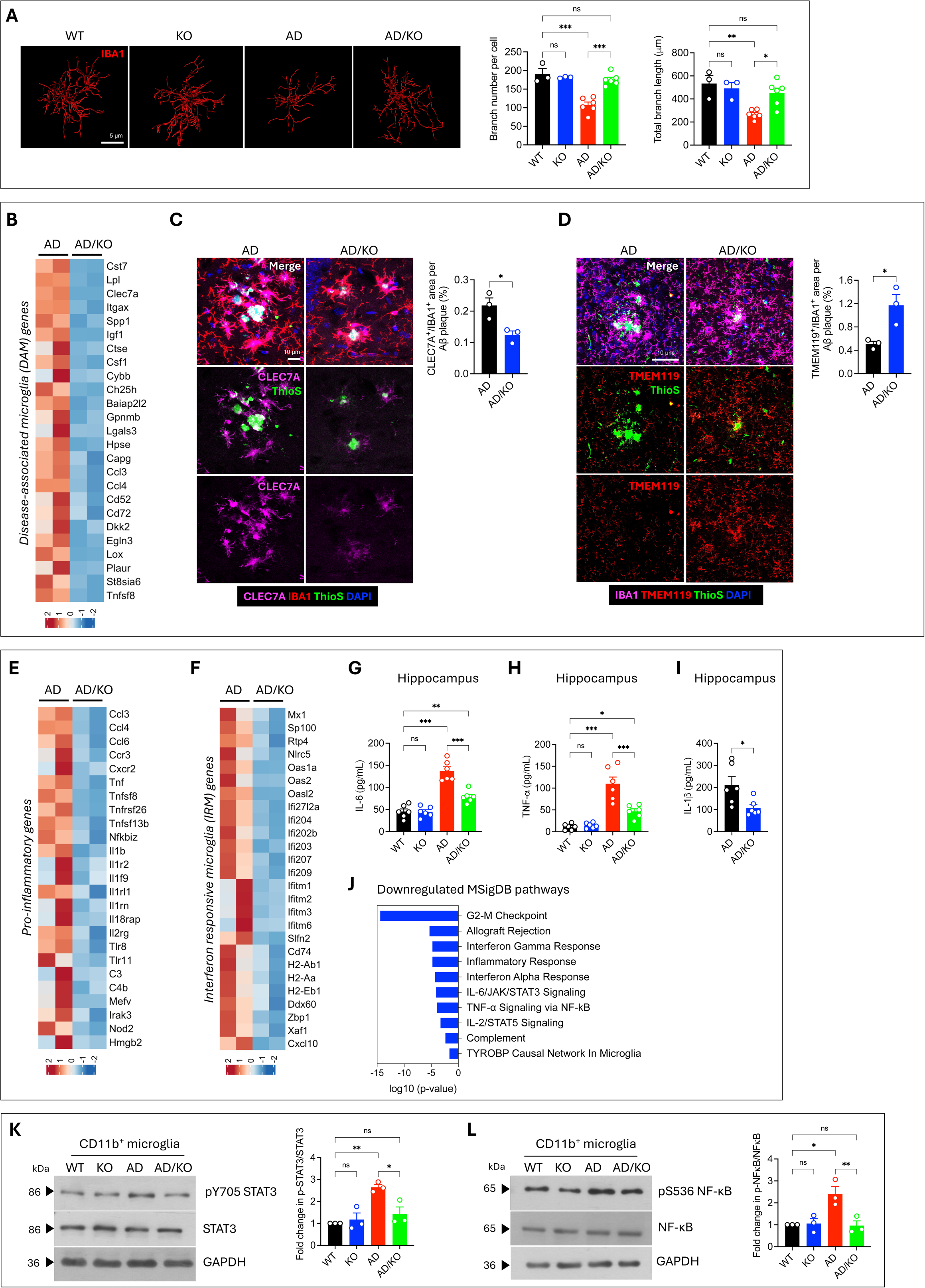
Asrij promotes inflammatory disease-associated microglia (DAM) activation state. (A) Representative renderings of the microglial skeleton using the ‘filament’ module of Imaris. Graphs show quantification of branch number per microglia and total branch length (n = 3 for WT and KO, n = 6 for AD and AD/KO). (B) Heatmap of significantly downregulated (*P* < 0.05, log_2_FC < −0.5) genes belonging to DAM signature. (C) Representative confocal images showing DAM marker, CLEC7A (magenta) on IBA1^+^ microglia (red) associated with ThioS^+^ plaques (green). Graph shows quantification of percent CLEC7A^+^ area normalized to IBA1^+^ area per plaque (n = 3). (D) Representative confocal images showing homeostatic marker, TMEM119 (red) on IBA1^+^ microglia (magenta) associated with ThioS^+^ plaques (green). Graph shows quantification of percent TMEM119^+^ area normalized to IBA1^+^ area per plaque (n = 3). (E, F) Heatmaps of significantly downregulated (*P* < 0.05, log_2_FC < −0.5) pro-inflammatory genes and IRM signature genes. (G, H, I) Graphs show quantification of IL-6, TNF-α and IL-1β levels (pg/mL) in the hippocampus by ELISA (n = 6). (J) Bar graph shows the top 10 significantly downregulated MSigDB hallmark pathways (*P* < 0.05). (K) Immunoblot analysis of phospho-STAT3 (tyrosine 705) and STAT3 levels in CD11b^+^ microglia. GAPDH is loading control. Graph shows quantification of fold change in p-STAT3 normalized to total STAT3 (n = 3). (L) Immunoblot analysis of phospho-NF-κB (serine 563) and NF-κB levels in CD11b^+^ microglia. GAPDH is loading control. Graph shows quantification of fold change in p-NF-κB normalized to total NF-κB (n = 3). Statistical significance between experimental groups was calculated by one-way ANOVA with Tukey’s post-hoc test (A, G-H and K-L), unpaired two-tailed Student’s t-test (C-D and I), and Fischer exact test (J). Error bars denote mean ± SEM. ns-non-significant, \**P* < 0.05, \*\**P* < 0.01, and \*\*\**P* < 0.001.

Single-cell RNA profiling of AD mouse models has identified that microglia adopt a unique transcriptional profile to become disease-associated microglia (DAM), also referred to as microglial neurodegenerative phenotype (MGnD). This state is characterized by the downregulation of homeostatic markers and upregulation of genes involved in innate immune signaling, phagocytosis, lipid metabolism, and inflammation (22,35,36). The downregulation of innate immune processes observed in our transcriptomic analysis spurred our interest to elucidate whether Asrij affects the acquisition of DAM transcriptional state in response to Aβ. Interestingly, among the most significantly downregulated were key genes involved in DAM phenotype, such as *Cst7*, *Lpl*, *Clec7a*, *Itgax*, *Csf1*, and *Ccl6* among others (Fig. 5B). We validated this block in DAM acquisition by evaluating levels of canonical DAM marker, C-Type lectin domain containing 7A (CLEC7A), on IBA1+ microglia surrounding Aβ plaques. AD/KO microglia expressed lower levels of CLEC7A than AD microglia (Fig. 5C). Also, DAM activation is accompanied by a reduction in the signature microglial homeostatic marker, transmembrane protein 119 (TMEM119) (35). We found that plaque-proximal AD/KO microglia showed higher TMEM119 levels than those in AD mice, suggesting their retention in a more homeostatic state (Fig. 5D). Altogether, our transcriptomic and imaging results highlight a critical role of Asrij in the transformation of homeostatic microglia to DAM in response to Aβ.

DAM can mount a pro-inflammatory or anti-inflammatory response depending on the disease stage (37, 38, 39). Inhibition of pro-inflammatory DAM response has been shown to reduce Aβ deposition and associated neuropathology in AD models (40–43). Hence, we characterized inflammatory changes that may accompany the attenuated DAM state seen in AD/KO mice. The majority of the downregulated genes in AD/KO microglia consisted of pro-inflammatory chemokines and cytokines such as *Ccl4*, *Tnf*, *Il1b*, immune receptors, and associated accessory proteins such as *Tnfsf8*, *Tnfrsf26*, *Il1r2*, *Il2rg*, pathogen recognition receptors including *Tlr8*, *Tlr11* and *Nod2* and complement factors, *C3* and *C4b* (Fig. 5E). Also, it was noteworthy, that 27 downregulated genes belong to the interferon response microglia (IRM) signature (44), such as the RNA recognition and binding factors, *Mx1*, *Oas2*, *Oas1a*, *Zbp1* and *Ddx60*, major histocompatibility complex (MHC) class II presentation such as *Cd74*, *H2-Ab1*, *H2-Aa*, and *H2-Eb1*, interferon-stimulated genes (ISGs) such as *Ifi27l2a*, *Ifi204*, *Ifitm1*, *Ifitm6* and *Cxcl10* (Fig. 5F). To validate RNA-seq results, we measured the concentration of pro-inflammatory cytokines, interleukin-6 (IL-6), tumor necrosis factor α (TNF-α) and interleukin-1β (IL-1β) in the brain homogenates using ELISA. While Asrij depletion alone in the absence of Aβ pathology does not affect cytokine levels, AD/KO mice brains had substantially reduced levels of IL-6, TNF-α and IL-1β as compared to AD mice (Fig. 5G-I). To identify which pathways are responsible for reduced inflammatory activation in Asrij deficient microglia, we performed pathway enrichment analysis on the downregulated DEGs using the molecular signatures database (MSigDB) hallmark gene set collection. This uncovered that interferon (IFN) gamma and alpha response, IL6/JAK/STAT3 signaling, TNF-α/NF-κB signaling, and IL-2/STAT5 signaling were significantly downregulated in AD/KO microglia (Fig. 5J). To validate reduced inflammatory pathway activation, we carried out immunoblotting analyses on MACS-sorted CD11b^+^ microglia. AD/KO microglia showed reduced levels of phosphorylated STAT3 and NF-κB, confirming the inhibition of inflammatory signaling (Fig. 5K-L). Taken together, these studies clearly demonstrate that Asrij depletion induces a shift to a less immunogenic and dampened pro-inflammatory microglial activation profile.

## Discussion

Alzheimer’s disease (AD) is a complex, multifactorial and progressive disease. An increase in Asrij levels in the brain positively correlates with the severity of AD pathology in mice and humans (4). Previous reports showed that primary neurons that are exposed to Aβ *in vitro*, upregulate Asrij and undergo mitochondria-mediated apoptosis, whereas Asrij depletion from neurons reduced Aβ-induced apoptosis (4). Hence, it is tempting to speculate a neuronal role for Asrij in neurodegeneration in AD patients. However, the role of Asrij in glial activation and neuroinflammation, which are among the most critical factors that modify the disease outcome, is not known. This necessitates a comprehensive analysis of the role of Asrij on AD pathology and progression, in an *in vivo* setting, that faithfully recapitulates the nuances of Aβ pathology. Therefore, we asked whether reducing Asrij levels *in vivo* is sufficient to mitigate the disease. Here, we show that Asrij depletion in the *APP/PS1* mice restores cognitive functions, reduces Aβ burden and neuronal damage, and decreases reactive astrogliosis. Mechanistically, Asrij suppresses mitochondrial activity and promotes neuroinflammatory signaling in AD microglia. Thus, increased Asrij in AD leads to microglial dysregulation and exacerbates Aβ pathology and cognitive dysfunction in mice.

Asrij regulates myeloid cell homeostasis and innate immune responses in *Drosophila* and mice. Further, Asrij is upregulated in AD microglia, suggesting that it has a role in microglial responses to Aβ pathology. We demonstrate that Asrij deletion significantly dampens microglial activation, as seen by reduced microgliosis and lesser microglial clustering around plaques. From the available reports, it is unclear whether microglial activation protects from AD progression or is detrimental of both. Activated microglia are necessary for Aβ phagocytosis and clearance, but can also propagate neuronal death and inflammation, via the production of cytokines (45). Also, complement-mediated aberrant synaptic pruning by activated microglia, contributes to excessive synaptic loss in AD (18). Microglia-driven neuroinflammation can further accelerate Aβ deposition (46). Given that AD/KO mice exhibit reduced Aβ plaques, it appears that reduced microglial activation and plaque association may be protective in this context, as opposed to other cases such as loss of TREM2, SYK, or VPS35, which leads to increased plaque burden and worsens AD pathology (15,29,33).

Asrij depletion leads to diminished microglial Aβ uptake and phagocytosis activation, with a concomitant reduction of Aβ burden. Though this finding may seem counter-intuitive, it is not surprising, given the growing evidence of poor correlation between microglial phagocytosis and plaque load (47). Recent studies have shown that microglial phagocytosis may, in fact, act to ‘build’ plaques rather than dismantle them. For example, the deletion of phagocytic receptors, Axl and Mer, in microglia, leads to an overall reduction of dense core plaques (24). Genetic or pharmacological depletion of microglia reduces plaque deposition (48–50). On the other hand, CARD9 deletion increases microgliosis as well as Aβ burden (14). Microglia also help seed and spread plaques, thus contributing to AD pathology (25,51). We found that Asrij deficient AD mice showed no change in plaque burden at 5 months of age, but had diminished Aβ plaque load and reduced plaque size from 10 months of age. Smaller Aβ aggregates act as seeds that spread and form larger Aβ plaques. Thus, it is tempting to speculate that Asrij may promote plaque seeding, deposition, and Aβ spread. Notably, although Asrij deficiency reduces microglial clustering around plaques, we noted reduced plaque volume and increased compaction. This suggests that microglial barrier function may be enhanced in AD/KO mice, even though fewer microglia are recruited to plaques. Therefore, Asrij has a critical role in influencing microglial barrier function and physical compaction of Aβ.

Plaque-associated microglia must maintain metabolic fitness, by upregulating energy metabolism and cellular biosynthetic pathways, to counter AD pathology (52,53). The extent of microglial metabolic dysfunction is a critical determinant of neuroinflammatory responses in AD (54,55). Our RNA-seq analysis reveals that Asrij depletion induces a unique transcriptional shift in AD microglia, marked predominantly by the upregulation of mitochondrial metabolism genes. Increased mitochondrial activity is particularly interesting, in light of previous studies linking Asrij to mitochondrial homeostasis. Asrij, an inner mitochondrial membrane protein, regulates mitochondrial dynamics and energy metabolism (56,57). It interacts with mitochondrial ETC complexes and negatively regulates Complex I activity, thus restraining oxidative phosphorylation (7). We hypothesize that Asrij depletion may alleviate this repression of mitochondrial activity and provide the energy boost required to maintain microglial metabolic fitness. However, the molecular mechanism by which Asrij controls microglial mitochondrial homeostasis warrants further investigation.

Single-cell RNA sequencing studies have uncovered that homeostatic microglia undergo a transcriptional shift to become DAM/MGnD during neurodegenerative diseases, chronic inflammation and aging (58). Our transcriptome analysis shows that Asrij facilitates the transition to the DAM state, as DAM signature genes and signaling pathways are dramatically downregulated in AD/KO mice. DAM are believed to be neuroprotective owing to their enhanced phagocytic and clearance functions (58). However, studies in AD mouse models have yielded confounding results, with reduced DAM resulting in either enhanced or reduced neuropathology. This may be because, under conditions of chronic inflammation, DAM get dysregulated and can be pro-inflammatory, anti-inflammatory, or a combination of both (59). For example, deletion of RIPK3, which reduces microglial pro-inflammatory activation, is accompanied by attenuated DAM and reduced Aβ pathology (40,41). Similarly, microglial cGAS deletion ameliorated neuroinflammation by inhibiting DAM emergence (42). Our study shows that Asrij is critical for the development of a pro-inflammatory DAM state. This is explained by the downregulation of several signaling pathways that contribute to inflammatory microglial activation. It is compelling that Asrij deficiency inhibited the interferon response microglia (IRM) profile, as observed by the remarkable reduction of IFN signaling genes. Hyperactive IFN signaling enhances neuroinflammation, synapse loss, and cognitive decline in AD (43). We also found reduced levels of pro-inflammatory cytokines in the brain milieu of AD/KO mice, suggesting reduced neuroinflammation. As neuroinflammation has been shown to aggravate Aβ pathology and cognitive impairments, it is reasonable to conclude that Asrij-dependent inflammatory microglial activation may contribute to AD progression. Mechanistically, Asrij acts as a scaffold protein to regulate multiple signaling pathways essential for immune homeostasis in *Drosophila* and mice (56). A similar function of Asrij, may be adapted in AD to fine-tune the immune signaling required for microglial activation. Given that our RNA-seq analysis revealed dramatic neuroinflammatory changes in microglia, we believe that the effects on AD pathology are, at least in part, driven by microglia-intrinsic functions, which may be confirmed by microglia-specific Asrij depletion. While Asrij knockout before disease onset retards AD progression, it will be interesting to see whether reducing Asrij levels after disease onset, may retard or ameliorate AD phenotypes.

Multiple studies have demonstrated the complexity of AD pathology, which depends on the gene affected, cell type and disease context. Several germline mutations in microglia-enriched genes that are implicated in AD may also affect peripheral immune cells. While modulation of microglial activation could be a promising therapeutic strategy, as many of the AD-related microglial receptors and signaling pathways are shared by peripheral myeloid cells, this could lead to undesirable effects (46). For example, the role of TREM2 in Aβ pathology is unclear. While one report indicates that TREM2 knockout reduces inflammatory macrophages and ameliorates Aβ pathology (60) another demonstrates that TREM2 knockout reduces microglial activation and increases Aβ pathology (61). Similarly, inhibiting SHIP-1 signaling leads to distinct effects depending on the disease stage and whether targeting is microglia-specific or pan-myeloid (62). These seemingly conflicting results highlight the importance of studying the role of molecular regulators in microglia, as well as in peripheral immune cells for safe and effective microglia-targeted AD therapies. Our study emphasizes that a tight balance of Asrij levels is essential for the coordination of its multiple roles. While increased Asrij levels promote AD, reducing Asrij levels in non-AD mice promotes uncontrolled myeloproliferation in the bone marrow. Here, we show that though Asrij maintains myeloid cell and mitochondrial homeostasis, it has a divergent role of immune activation in microglia. Therefore, our study uncovers a novel mechanism of Asrij-dependent microglial dysfunction that drives neuroinflammatory microglial responses. Given the diverse functions of Asrij, therapeutic modulation of its levels in AD will likely result in distinct clinical outcomes, based on the disease stage and systemic responses. Our study gives insights into microglial biology that could help devise strategies to prevent or retard AD progression and improve clinical outcomes.

## Conclusions

In summary, we show that Asrij depletion ameliorates Aβ pathology, neuronal and synaptic damage and gliosis, and improves behavioral performance in *APP/PS1* mice. Asrij suppresses microglial mitochondrial activity and promotes the development of DAM, which likely leads to worsening of AD pathology with increase in Asrij levels. Depleting Asrij in an AD context limits neuroinflammatory signaling activation in microglia, enabling neuroprotective microglial responses. Collectively, our reducing Asrij may help retard AD. Our work positions Asrij as a critical molecular regulator that links microglial dysfunction to AD pathogenesis.

## Abbreviations

AD: Alzheimer’s disease
Aβ: Amyloid beta
GWAS: Genome-wide association studies
OCIAD1: Ovarian Carcinoma Immunoreactive Antigen Domain protein 1
GSK3β: Glycogen synthase kinase 3 beta
DAM: Disease-associated microglia
WT: Wild type
KO: Knockout
PCR: Polymerase chain reaction
JNCASR: Jawaharlal Nehru Centre for Advanced Scientific Research
OFT: Open field test
MWM: Morris water maze
PBS: Phosphate-buffered saline
PFA: Paraformaldehyde
RNA: Ribonucleic acid
RT: Room temperature
FBS: Fetal bovine serum
DAPI: 4′,6-diamidino-2-phenylindole
HEPES: 4-(2-hydroxyethyl)-1-piperazineethanesulfonic acid
EDTA: Ethylenediaminetetraacetic acid
DTT: Dithiothreitol
PMSF: Phenylmethylsulfonyl fluoride
SDS: Sodium dodecyl sulfate
HRP: Horseradish peroxidase
ECL: Clarity western enhanced chemiluminescence
HBSS: Hanks’ balanced salt solution
BSA: Bovine serum albumin
DPBS: Dulbecco’s phosphate-buffered saline
MACS: Magnetic-activated cell sorting
FACS: Fluorescence-activated cell sorting
TMRM: Tetramethylrhodamine, Methyl Ester, Perchlorate
FSC-A: Forward scatter area
SSC-A: Side scatter area
FSC-H: Forward scatter height
SSC-W: Side scatter width
MFI: Median fluorescence intensity
IMDM: Iscove’s modified Dulbecco’s medium
ELISA: Enzyme-linked immunosorbent assay
RT-qPCR: Reverse transcription-quantitative polymerase chain reaction
QC: Quality control
rRNA: ribosomal RNA
PCA: Principal component analysis
MSigDB: Molecular signatures database
SEM: Standard error of mean
ANOVA: Analysis of variance
ThioS: Thioflavin S
APP: Amyloid precursor protein
CTF: Carboxy-terminal fragments
LAMP1: Lysosomal-associated membrane protein 1
PSD95: Post synaptic density 95
GFAP: Glial fibrillary acidic protein
S100B: S100 calcium-binding protein B
IBA1: Ionized calcium-binding adapter molecule 1
CD68: Cluster of differentiation 68
RNA-seq: RNA sequencing
GO: Gene ontology
DEG: Differentially expressed genes
TMRM: Tetramethylrhodamine, Methyl Ester, Perchlorate
ROS: Reactive oxygen species
mTOR: Mammalian target of rapamycin
MGnD: Microglial neurodegenerative phenotype
CLEC7A: C-Type lectin domain containing 7A
TMEM119: Transmembrane protein 119
IRM: Interferon response microglia
ISG: Interferon-stimulated genes
MHC: Major histocompatibility complex
IL-6: Interleukin-6
TNF-alpha: Tumor necrosis factor α
IL-1beta: Interleukin-1β
IFN: Interferon
KEGG: Kyoto encyclopedia of genes and genomes
GAPDH: Glyceraldehyde-3-phosphate dehydrogenase
STAT3: Signal transducer and activator of transcription 3
NF-κB: Nuclear factor kappa-light-chain-enhancer of activated B cells
ETC: Electron transport chain
BLiSC: Bangalore Life Science Cluster

## Declarations

### Ethics approval and consent to participate

All mice experiments were approved by the JNCASR institutional animal ethics committee (project #MSI012) and were conducted in accordance with the standard guidelines. This work did not involve the use of material from human subjects.

### Consent for publication

All authors have read and approved the final manuscript for publication.

### Availability of data and materials

All data supporting the findings of this study are available in the paper and its supplementary figures. All the original raw data and analysis files are available with the corresponding authors upon reasonable request. RNA sequencing data has been deposited into the gene expression omnibus (GEO) database under accession number GSE272625.

### Authors’ contributions

Conceptualization: P.D, T.G.R and M.S.I. Methodology, experiments, and data analysis: P.D and M.R. Data discussions: P.D, M.R., T.G.R and M.S.I. Writing - original draft: P.D. and M.S.I. Writing - review and editing: P.D, M.R., T.G.R, M.S.I. Funding acquisition and resources: M.S.I and T.G.R.

## Acknowledgments

The authors thank members of Inamdar laboratory for valuable suggestions; Aishwarya Prakash for RNA-seq data analysis; Dr. Ashish Kumar for help with mice maintenance; Hiyaa Ghosh lab at the National Centre for Biological Sciences (NCBS), Bangalore for help with reagents and protocols; Bioimaging, flow cytometry and animal facilities at JNCASR. Graphical abstract and schematics were prepared using BioRender.

## Supplementary figure legends

**Figure S1.**
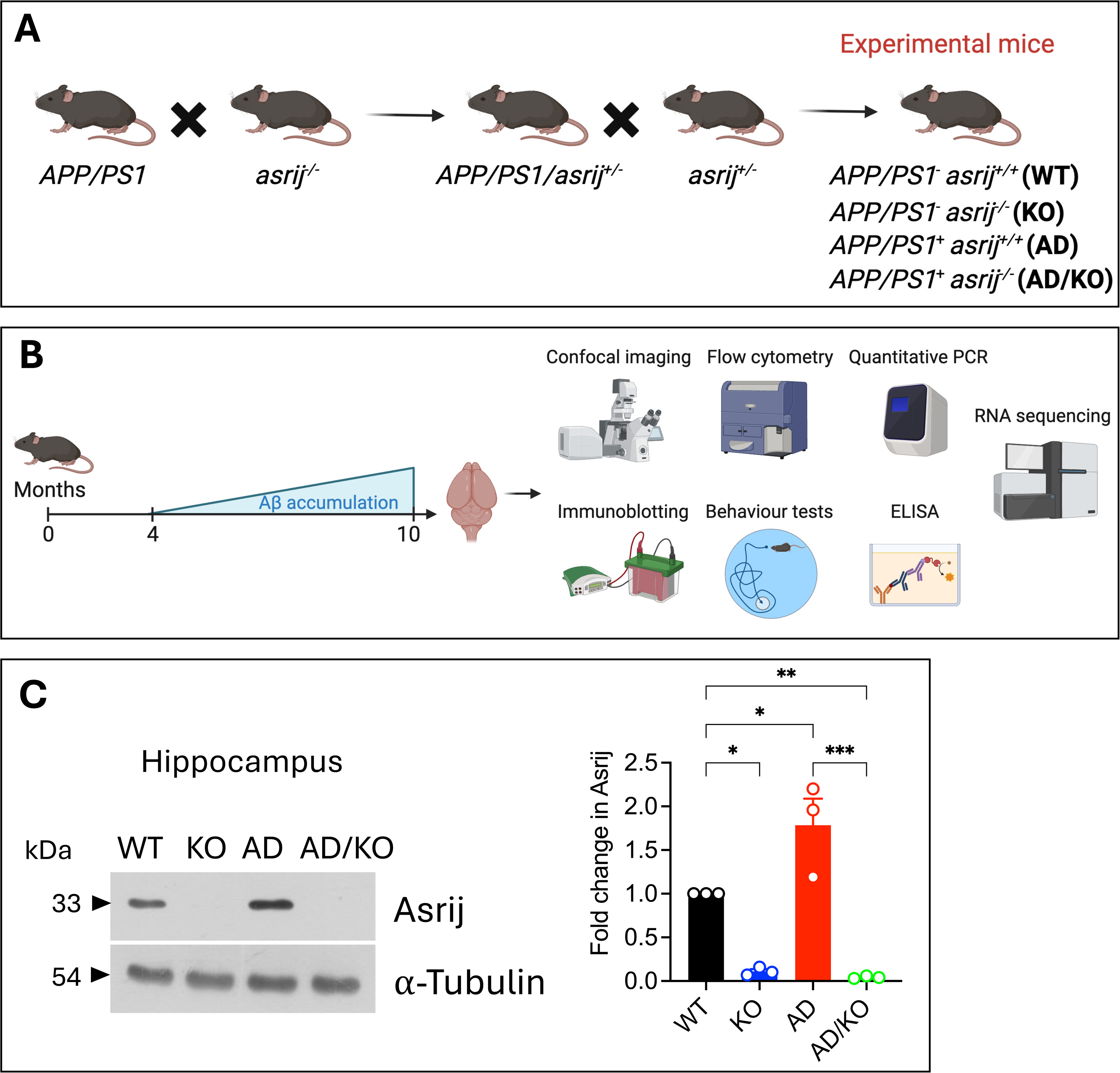
Generation, characterization, and validation of *APP*/*PS1*/*asrij*^−/−^ mice. (A) Schematic representation of breeding strategy to obtain homozygous deletion of *asrij* in *APP/PS1* mice. (B) Schematic illustrating experimental design for characterization of Aβ pathology in mice. (C) Immunoblot analysis of Asrij levels in the hippocampus. α-tubulin is loading control. Graph shows fold change in Asrij levels normalized to α-tubulin (n = 3). Statistical significance between experimental groups was calculated by one-way ANOVA with Tukey’s post hoc test. Error bars denote mean ± SEM. ns-non-significant, \**P* < 0.05, \*\**P* < 0.01, and \*\*\**P* < 0.001.

**Figure S2.**
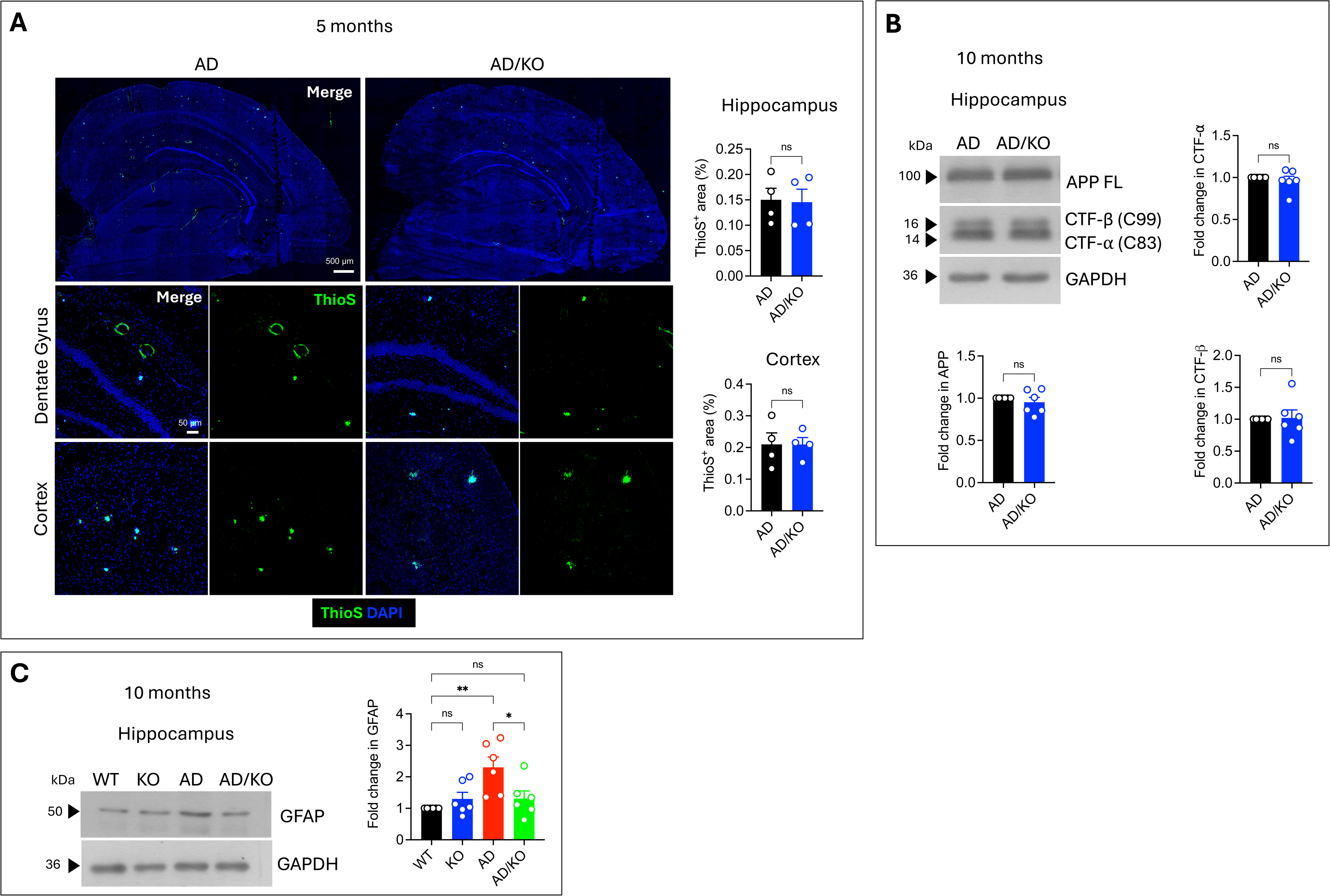
Asrij depletion does not affect Aβ plaque accumulation at 5 months and APP processing at 10 months of age. (A) Representative confocal images of brain sections showing ThioS staining (green) for Aβ plaques. DAPI (blue) marks the nuclei. Graphs show quantification of the percent area covered by ThioS^+^ staining in the hippocampus and cortex (n = 4). (B) Immunoblot analysis of full-length amyloid precursor protein (APP) and carboxyl-terminal fragments (CTF), CTF-α and CTF-β in the hippocampus. GAPDH is loading control. Graphs show quantification of fold change in APP, CTF-α, and CTF-β levels normalized to GAPDH (n = 6). (C) Immunoblot analysis of GFAP levels in the hippocampus. GAPDH is loading control. Graph shows quantification of fold change in GFAP levels normalized to GAPDH (n = 6). Statistical significance between experimental groups was calculated by unpaired two-tailed Student’s t-test (A-B) and one-way ANOVA with Bonferroni’s post hoc test (C). Error bars denote mean ± SEM. ns-non-significant, \**P* < 0.05, \*\**P* < 0.01, and \*\*\**P* < 0.001.

**Figure S3.**
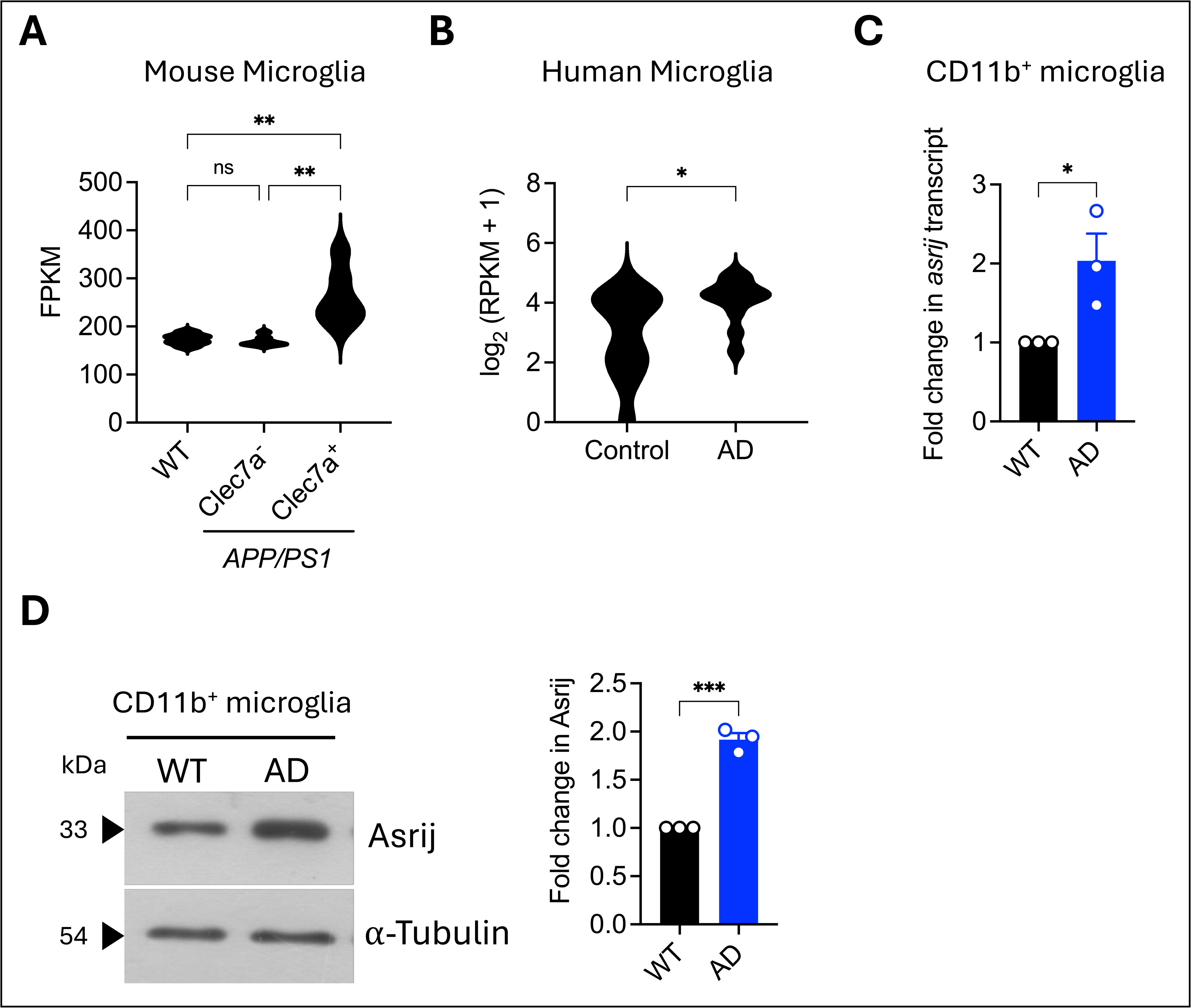
Asrij transcript and protein levels are increased in AD microglia. (A) Violin plot showing *asrij* transcript expression in mouse Clec7a^−^ (non-plaque-associated) and Clec7a^+^ (plaque-associated) microglia (n = 6). Data plotted from Krasemann *et al*., 2017. (B) Violin plot showing *asrij* transcript expression in human microglia (n = 11). Data plotted from Mathys, Davila-Velderrain *et al*., 2019. (C) RT-qPCR analysis of *asrij* expression in MACS-sorted CD11b^+^ mouse microglia. Graph shows quantification of fold change in *asrij* transcript normalized to β-actin expression (n = 3). (D) Immunoblot analysis of Asrij protein levels in MACS-sorted CD11b^+^ mouse microglia. Graph shows quantification of fold change in Asrij levels normalized to α-tubulin (n = 3). Statistical significance between experimental groups was calculated by one-way ANOVA with Bonferroni’s post hoc test (A) and unpaired two-tailed Student’s t-test (B-D). Error bars denote mean ± SEM. ns-non-significant, \**P* < 0.05, \*\**P* < 0.01, and \*\*\**P* < 0.001.

**Figure S4.**
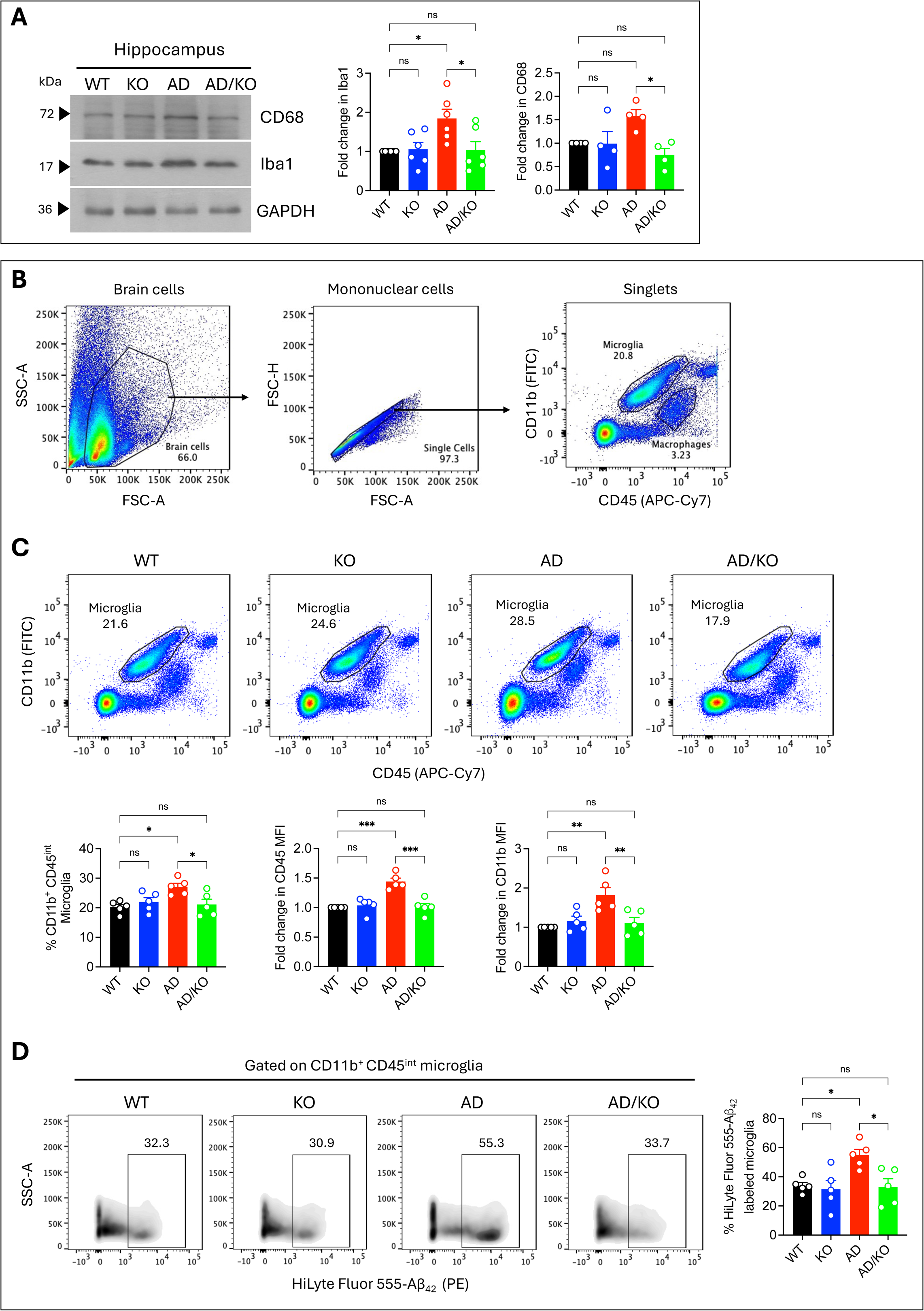
Reduced microglial phagocytic activation in Asrij deficient *APP/PS1* mice. (A) Immunoblot analysis of IBA1 and CD68 levels in the hippocampus. GAPDH is loading control. Graphs show quantification of fold change in IBA1 (n = 6) and CD68 (n = 4) levels normalized to GAPDH. (B) Gating strategy for FACS analysis of microglia. (C) Representative flow cytometry dot plots showing the frequency of microglia. Graph shows quantification of the percentage of CD11b^+^ CD45^int^ microglia (n = 5). Graphs show quantification of fold change in median fluorescence intensity (MFI) of CD45 and CD11b in microglia (n = 5). (D) Representative flow cytometry density plots showing the frequency of microglia positive for HiLyte Fluor 568 labeled Aβ_42_ oligomers. Graph shows quantification of the percentage of HiLyte Fluor 568-Aβ_42_ stained microglia. Statistical significance between experimental groups was calculated by one-way ANOVA with Bonferroni’s post hoc test. Error bars denote mean ± SEM. ns-non-significant, \**P* < 0.05, \*\**P* < 0.01, and \*\*\**P* < 0.001.

**Figure S5.**
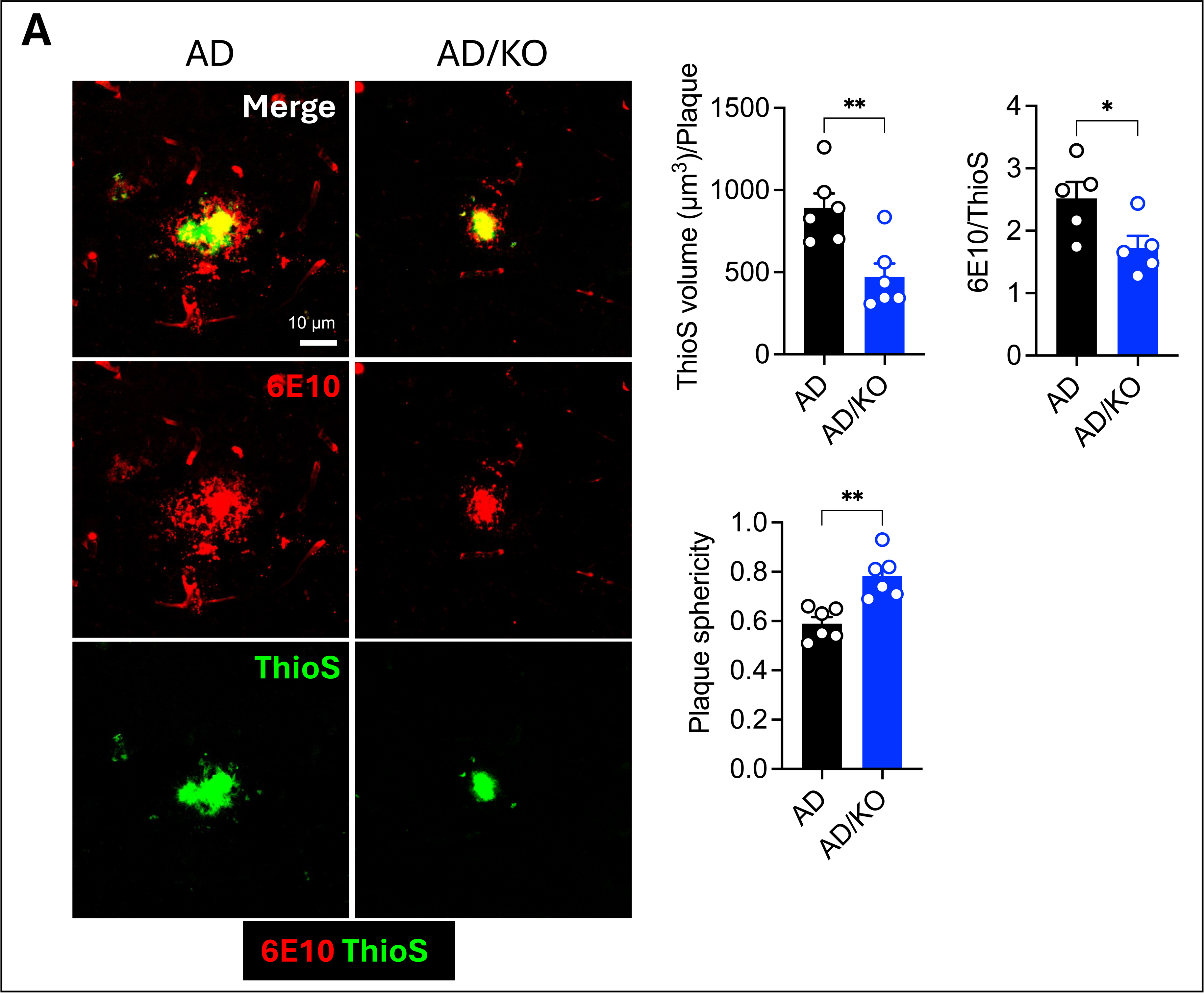
Asrij depletion promotes Aβ plaque compaction in *APP/PS1* mice. (A) Representative confocal images showing 6E10^+^ fibrillar plaque (red) and ThioS^+^ compact plaque (green). Graphs show quantification of ThioS^+^ volume per plaque (μm^3^), plaque sphericity, and the ratio of 6E10 and ThioS area (n = 6). Statistical significance between experimental groups was calculated by unpaired two-tailed Student’s t-test. Error bars denote mean ± SEM. ns-non-significant, \**P* < 0.05, \*\**P* < 0.01, and \*\*\**P* < 0.001.

**Figure S6.**
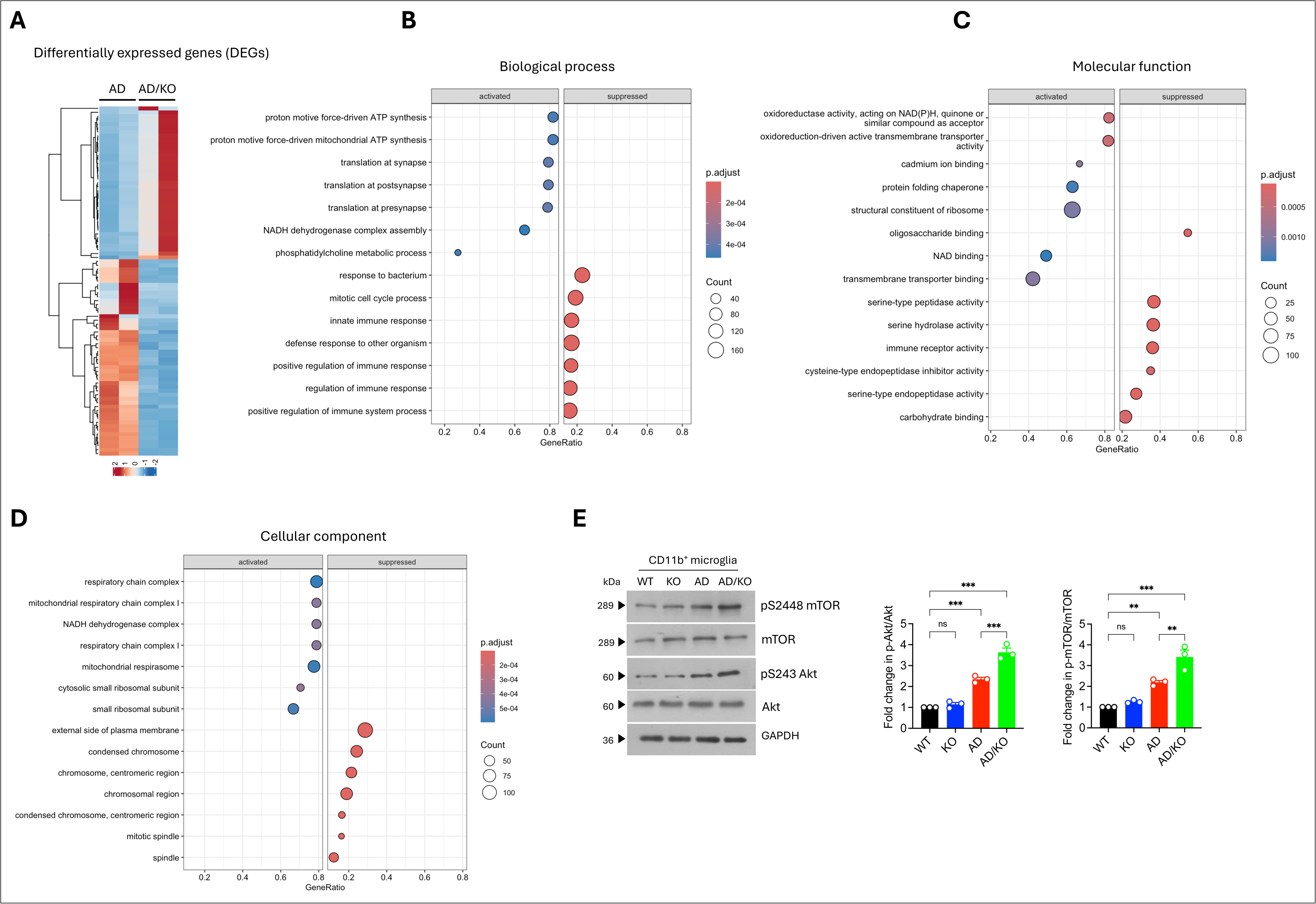
Upregulation of metabolic processes and downregulation of immune processes in Asrij deficient *APP/PS1* mice. (A) Heatmap of all significantly affected DEGs (*P* < 0.05). (B-D) Enrichment dot plots showing the top 7 significantly activated and repressed GO terms (adjusted *P* < 0.05) using clusterProfiler R package. (E) Immunoblot analysis of phospho-mTOR (serine 2448), mTOR, phospho-Akt (serine 243), and Akt. GAPDH is loading control. Graphs show fold change in p-mTOR normalized to total mTOR and fold change in p-Akt normalized to total Akt. Statistical significance between experimental groups was calculated by two-sided Wald test (A), hypergeometric test (B-D), and one-way ANOVA with Tukey’s post-hoc test (E). Error bars denote mean ± SEM. ns-non-significant, \**P* < 0.05, \*\**P* < 0.01, and \*\*\**P* < 0.001.

## Supplementary table

**Supplementary table:** A list of differentially expressed genes in *APP/PS1*^+^/*asrij*^−/−^ (AD/KO) CD11b^+^ microglia using RNA sequencing.

